# Discovery and performance of DNA methylation panels for cancer detection and classification in blood

**DOI:** 10.1101/2025.09.16.676542

**Authors:** Gennady Margolin, Sara R. Bang-Christensen, Sara A. Grimm, Brian D. Bennett, Peter Kilfeather, Kevin Fedkenheuer, Sabina Pathan, Nichelle C. Whitlock, Peter A. Pinto, Leszek J. Klimczak, S. Katie Farney, Nader Jameel, Yun-Ching Chen, Hanna M. Petrykowska, Adam G. Sowalsky, Jian-Liang Li, Laura Elnitski

**Author notes:** Co-first author (Gennady Margolin, Sara R. Bang-Christensen). **Correspondence: L. Elnitski**, The National Human Genome Research Institute, National Institutes of Health, 20892 Bethesda, MD, USA.

## Abstract

Examining DNA in a liquid biopsy for non-invasive cancer detection relies on identifying dilute signal in a high background. This study aims to identify DNA methylation biomarkers for multi-cancer detection. Utilizing large tissue datasets, we apply novel search algorithms to discover confined biomarker panels capable of distinguishing tumor from normal and determining the tissue of origin. We explore the applicability to blood-based testing using targeted methylation sequencing followed by machine learning classification. We present an 8-marker panel, which successfully predicts tumors across 14 types with a 91% average sensitivity, maintaining a low false positive rate (< 0.04%). Additionally, a panel of 39 CpG sites exhibits accuracies ranging from 69% to 98% for identifying tissue of origin. When tested on 114 patient plasma samples (colon, liver, pancreatic, prostate, and stomach cancer), the 8-marker panel obtains an AUC of 0.78 with a 78% sensitivity among 32 early-stage patients (stage I-II), and 60% overall. Using the 39-marker panel in a multi-class classification model selecting only the best match, 54% of tumor samples were on average correctly assigned to the tissue of origin, and up to 80% when allowing more inclusive criteria. Using a limited set of biomarkers, our work contributes to advancing non-invasive cancer diagnostics.

## 1 Introduction

Liquid biopsies, which are body fluid-based tests carrying information about solid tumors, have emerged as a minimally invasive and cost-effective approach to tumor detection. As a testing modality, liquid biopsy is expected to provide a sensitive means of disease detection that complements the use of existing imaging and biopsy approaches. In practice, the ability to repeatedly sample a patient via liquid biopsy enables frequent assessment that is safer than repeated tissue biopsy [1], can be more sensitive than imaging [2], and enables comparisons of biomarker levels before and after surgery or therapeutic intervention. Compared to conventional protein glycosylation biomarkers (e.g., serum prostate-specific antigen, CA15, carcinoembryonic antigen [3]), which are not specific for neoplasia, circulating tumor DNA (ctDNA) can achieve greater sensitivity and specificity when used to detect cancer. ctDNA can also improve testing outcomes when combined with conventional markers [4–6]. Moreover, liquid biopsy represents a direct implementation of precision medicine, as it can guide treatment based on disease markers specific to an individual patient. Once known, these markers can be used for monitoring disease [7, 8] and detecting residual disease [9], therapeutic resistance [10, 11], and disease recurrence [12].

DNA mutations have demonstrated the feasibility of detecting ctDNA from circulating cell-free DNA (cfDNA). For example, Bettegowda et al. [13] detected known DNA mutations in ctDNA from blood samples collected from patients with 15 cancer types, including early-stage disease. Strategies that rely on mutation detection perform best with prior knowledge of the mutations present in patient tumors; enabling bespoke detection tests to look for those sequence changes in blood samples [14, 15]. A downside to using mutations to detect cancer is the inability to know which mutations to test without *a priori* knowledge, particularly given the low frequency of most cancer driver mutations within any cancer type, but especially across cancer types [16]. This ambiguity necessitates whole genome sequencing to discover mutations that are likely cancerous.

In contrast to mutations, DNA methylation alters the epigenome of tumors, so they are distinct from healthy tissues. These alterations are stable over time, recur with high frequency in specific tumor types, and can be used to reliably detect tumors from different organs [17, 18]. For example, our lab previously identified a DNA methylation marker in the *ZNF154* promoter as a putative pan-cancer biomarker [19, 20]. Most recently, we showed that this marker can identify the presence of ovarian, pancreatic, liver and colon cancer when used in a sensitive detection assay on patient plasma samples and that this assay can complement and improve results obtained with the standard plasma protein markers or cancer-specific mutations [6, 21].

In addition to binary assessments evaluating presence or absence of cancer, several methods have been proposed to identify sites of tumor origins from blood-based testing, utilizing varying strategies and analytes. These methods often interrogate an extensive number of target sites, including whole genome bisulfite sequencing (WGBS), to look for cancer type-specific alterations in DNA methylation. Such alterations can be determined *in silico* on a broad scale, as illustrated by 2,880 genomic locations in methylation haplotype blocks [22], collections of 1,250 and 584 hypermethylated and hypomethylated markers, respectively [23], and 14,716 markers defined by immunoprecipitating cell-free methylated DNA, offering a bisulfite-free approach [24]. Lab-based testing approaches that include clinical data are exemplified by CANCER SEEK, which combines protein markers with DNA mutations to identify 8 cancer types [25]. Proprietary approaches have been reported by the company GRAIL, which employ targeted methylation sequencing of cfDNA [26]. One recent study by GRAIL reported 43% sensitivity and 99.5% specificity in a test of 6,662 individuals using a panel of roughly 1 million CpG sites to differentiate up to 50 cancer types [27]. Additional advances in blood-based testing are accruing quickly. Although promising, however, these approaches are potentially dependent on extensive sequencing of cfDNA from plasma, which is costly and may prohibit widespread implementation. As such, some detection methods are unlikely to translate readily to small, remote clinics, underserved populations, or uninsured individuals.

We reasoned that a straightforward blood-based test capable of identifying and distinguishing between multiple cancer types requires a panel of methylation markers that reliably identifies the source of a tumor, has a small number of relevant sites, and can be implemented without whole genome sequencing. To this end, here we analyze The Cancer Genome Atlas (TCGA) methylation array data (Illumina Infinium BeadChip) [28] and a collection of over 2,700 peripheral blood samples from donors without cancer [29] to identify candidate markers for such a test. We describe markers whose methylation can be used for not only binary assessment of tumor presence or absence, but also to discriminate between up to 13 different cancer types originating in 11 different tissues/organs thereby aiding in identifying the tissue of origin. We validate our findings using independent Illumina methylation array assessment and follow up by testing these marker panels in cfDNA from plasma and applying machine learning algorithms for classifying tumor and normal samples. In short, our study presents the discovery, validation and initial clinical testing of two novel, size-restricted DNA methylation marker panels capable of determining tumor presence and tissue of origin, providing the foundation for potential development into a clinical application.

## 2 Materials and Methods

### 2.1 TCGA methylation array data used for marker discovery and initial testing

To select markers/probes and analyze their performance, we used TCGA Infinium Human Methylation 450K BeadChip array data from 14 solid tumor types (Table 1): bladder urothelial carcinoma (BLCA), breast invasive carcinoma (BRCA), colon adenocarcinoma (COAD), head and neck squamous cell carcinoma (HNSC), kidney renal clear cell carcinoma (KIRC), kidney renal papillary cell carcinoma (KIRP), liver hepatocellular carcinoma (LIHC), lung adenocarcinoma (LUAD), lung squamous cell carcinoma (LUSC), pancreatic adenocarcinoma (PAAD), prostate adenocarcinoma (PRAD), rectum adenocarcinoma (READ), stomach adenocarcinoma (STAD), and uterine corpus endometrial carcinoma (UCEC). For each type of cancer, data from both tumor and normal tissue were available; we append an “.N” or “.T” to distinguish normal and tumor samples, respectively. In our analyses, we pooled colon adenocarcinoma (COAD) and rectum adenocarcinoma (READ) samples to form a new colorectal adenocarcinoma (CRAD) category, as the data were virtually indistinguishable in our initial analysis, as confirmed in TCGA studies [30, 31], explaining why we say we discriminate among 13 cancer types.

**Table 1.** Counts of samples used for the peripheral blood reference and TCGA tissue types.

| Blood reference or TCGA Cancer Type | Description | Normal (N) | Tumor (T) |
| --- | --- | --- | --- |
| reference | Peripheral blood | 2,711 | 0 |
| BLCA | Bladder urothelial carcinoma | 20 | 201 |
| BRCA | Breast invasive carcinoma | 96 | 676 |
| COAD | Colon adenocarcinoma | 38 | 274 |
| HNSC | Head and neck squamous cell carcinoma | 50 | 426 |
| KIRC | Kidney renal clear cell carcinoma | 160 | 296 |
| KIRP | Kidney renal papillary cell carcinoma | 45 | 156 |
| LIHC | Liver hepatocellular carcinoma | 50 | 151 |
| LUAD | Lung adenocarcinoma | 32 | 437 |
| LUSC | Lung squamous cell carcinoma | 42 | 359 |
| PAAD | Pancreatic adenocarcinoma | 9 | 65 |
| PRAD | Prostate adenocarcinoma | 49 | 248 |
| READ | Rectum adenocarcinoma | 7 | 96 |
| STAD | Stomach adenocarcinoma | 2 | 260 |
| UCEC | Uterine corpus endometrial carcinoma | 46 | 407 |

### 2.2 Blood reference methylation array data used for marker discovery and initial testing

Because our ultimate goal was to detect tumors using blood plasma samples, we used GSE55763 samples [29] from the Gene Expression Omnibus (GEO) repository [32] as references for healthy blood DNA methylation levels (Table 1). The GSE55763 dataset contains over 2,700 peripheral blood samples, with more than 1,600 samples from healthy subjects with no reported condition/pathology and more than 1,000 samples from individuals with type 2 diabetes. No information was available to determine which category (healthy vs type 2 diabetes) the samples belonged to. However, 95% of all markers had a methylation beta-value standard deviation < 0.07, indicating negligible to small differences between the two categories for the majority of markers. Given this observation, and the fact that in our marker selection algorithms we filter out markers with large standard deviations (see ‘Marker selection strategy’ below), we concluded that the presence of ∼40% of blood samples from individuals with type 2 diabetes in this dataset does not preclude its use as a source of peripheral blood reference samples for our purposes.

### 2.3 Independent validation methylation array data

To validate the performance of the probes we selected, we used the following GEO methylation array datasets as validation datasets: GSE37754, GSE49149, GSE53051, GSE55479, GSE61441, GSE66695, and GSE69914 (summarized in Table 2). We also used WGBS data from plasma generated by Chan et al. (2013) [33] and tissue and cell line samples from Vidal et al. 2017 [34]. Additionally, we used tissue and peripheral blood cell DNA from several non-cancer conditions, whose methylation array data served as negative controls (GSE32148, GSE85566, GSE50874, GSE49542, and GSE87621), as well as Dayeh et al. (2014) [35] (summarized in Table 3).

**Table 2.** Validation samples: Tumors and normal controls.

| Data Source | Description | Sample Count | Format |
| --- | --- | --- | --- |
| Chan et al. 2013 [33] | solid HCC | 15 | WGBS |
|  | normal plasma | 32 | WGBS |
|  | HCC plasma | 26 | WGBS |
| GSE49149 | pancreatic ductal adenocarcinoma | 167 | Infinium 450K array |
|  | adjacent non-tumor | 29 | Infinium 450K array |
| GSE37754 | breast tumor | 62 | Infinium 450K array |
|  | adjacent noncancerous tissue | 10 | Infinium 450K array |
| GSE66695 | breast tumor | 80 | Infinium 450K array |
|  | normal tissue | 40 | Infinium 450K array |
| GSE69914 | breast cancer | 305 | Infinium 450K array |
|  | cancer-brca1 | 3 | Infinium 450K array |
|  | normal | 50 | Infinium 450K array |
|  | normal adjacent | 42 | Infinium 450K array |
|  | normal-brca1 | 7 | Infinium 450K array |
| GSE61441 | renal clear cell carcinoma | 46 | Infinium 450K array |
|  | matched normal kidney tissue | 46 | Infinium 450K array |
| GSE53051 | breast cancer | 14 | Infinium 450K array |
|  | breast normal | 10 | Infinium 450K array |
|  | colon cancer | 35 | Infinium 450K array |
|  | colon normal | 18 | Infinium 450K array |
|  | lung cancer | 9 | Infinium 450K array |
|  | lung normal | 11 | Infinium 450K array |
|  | pancreas cancer | 29 | Infinium 450K array |
|  | pancreas normal | 12 | Infinium 450K array |
| GSE55479 | prostate cancer | 143 | Infinium 450K array |
| Vidal et al. 2017<br>[34] | 11 tumor and tumor cell line samples, 5 normal samples, and 1 B-lymphocyte sample | 17 | WGBS |

**Table 3.** Illumina methylation microarray data from non-cancer conditions and normal control samples.

| Data source | Description | Sample count | # Called as tumor | # Called as normal | Format |
| --- | --- | --- | --- | --- | --- |
| GSE32148 | peripheral blood cells: Crohn's' disease | 17 | 0 | 17 | Infinium 450K array |
|  | peripheral blood cells: ulcerative colitis | 11 | 0 | 11 | Infinium 450K array |
|  | peripheral blood cells: normal controls | 20 | 0 | 20 | Infinium 450K array |
| GSE85566 | airway epithelial cells: asthma | 74 | 0 | 74 | Infinium 450K array |
|  | airway epithelial cells: control | 41 | 0 | 41 | Infinium 450K array |
| GSE50874 | chronic kidney disease | ~20<sup>a</sup> | 0 | ~20<sup>a</sup> | Infinium 450K array |
|  | kidney control | ~65<sup>a</sup> | 0 | ~65<sup>a</sup> | Infinium 450K array |
| GSE49542 | non-alcoholic fatty liver disease [frozen liver biopsies] | 59 | 0 | 59 | Infinium 450K array |
| GSE87621 | endometriosis [cultured primary cells] | 4 | 0 | 4 | Infinium 450K array |
|  | control [cultured primary endometrial stromal cells] | 5 | 0 | 5 | Infinium 450K array |
| PMID: 24603685 [35] | pancreatic islets: T2D | 15 | 0 | 15 | Infinium 450K array |
|  | pancreatic islets: non-diabetic | 34 | 0 | 34 | Infinium 450K array |

<sup>a</sup>85 micro-dissected human kidney tubule epithelial samples without identification comprising: 20 samples from patients with diabetic *kidney* disease and 65 controls (mixed as 1/3 hypertension, 1/3 diabetes mellitus and 1/3 healthy).

### 2.4 Methylation analysis

To prepare the array data, Illumina Infinium array beta methylation values were normalized as previously published [19, 36]. We used the hg19 human reference genome and the Illumina annotation file for the Illumina Infinium HumanMethylation450 BeadChip array to obtain information on probe identifiers. Reads from whole-genome bisulfite sequencing (WGBS) data were quality-checked with FastQC, adapters trimmed with Picard (v. 0.14.4, MarkIlluminaAdapters to SamToFastq), and aligned with Bismark (v.0.14.4) [37] to the human reference genome (hg19). Alignments were coordinate-sorted and duplicate reads removed with Picard MarkDuplicates. Cytosine methylation was extracted with bismark_methylation_extractor. Downstream processing and visualization were performed in Perl and R.

### 2.5 Marker selection strategy

For the purpose of selecting markers that discriminate tumor DNA from the background of heathy tissue DNA, we determined probes whose methylation in normal reference samples represented the extremes of the scale, i.e., virtually absent or saturated, with minimal variability. At these loci, even a weakly abnormal signal becomes apparent against the background. To make a pool of candidate tumor-normal (T-N) markers, we required that the median methylation value of each marker differed substantially in at least one tumor type from the blood reference (see Supplemental Methods). In addition, we required that all normal tissue values were similar to the blood reference. Iterative cycles of this approach yielded a set of eight probes with capacity to discriminate tumor and normal samples such that each tumor type was detected by at least 2 probes. We call them “unconditional” T-N probes, because these probes can be used to differentiate tumors of all types within the collection of 13 tumor types we consider here, from normal samples and peripheral blood, without knowing the type *a priori*. To classify samples, methylation beta values at each marker were binarized with ‘1’ being a methylation value far from the blood reference and ‘0’ being similar to the reference (see Supplemental Methods). A sample was called as tumor if at least one of the binarized values was 1; otherwise, it was called as normal.

For the purpose of selecting markers that assign tumor samples into one of the TCGA types (or blood reference) we again required extreme methylation values in the reference samples with substantially different methylation levels in tumor samples summarized by type. Each probe had to represent at least one tumor type, whereas the probe’s methylation in all negative tumor types had to satisfy the same thresholds as were valid for the reference (Supplemental Methods). This approach yielded a pool of candidate markers, which through multiple iterations were selected based on an entropy measure resulting in several sets of probes successfully splitting all classification types. Similarly to the T-N call probes, the beta values were binarized based on extreme opposite (‘1’) or similar (‘0’) methylation levels compared to reference samples. The final panel of tissue classification probes contained 39 markers, yielding a binary profile of 39 values for each tumor type. To perform tumor type classification of individual samples, the beta values along the 39 markers were binarized and the mean Euclidean distance to each classification type was calculated. Thus, for each sample we had a vector of mean distances to the classification types (Figure S1). The simplest classification was assignment to the closest scoring tumor type, whereas alternate criteria were adopted to assess the reliability of each classification within an acceptable range. Thus, we additionally checked (1) whether the correct class ranked :< 2 (i.e., best or second-best match, with nuances in case of ties) and recorded the rank of the correct class, and (2) whether the correct class could be found within a specified range starting from the sample and extending a pre-specified distance into the Euclidean space, again recording the number of classes within that range (Supplemental Methods). Finally, we considered two ways to adjust for the classification bias due to different distributions of distances from individual samples to the correct classification type in different tumor types. For that we tested quantile fraction fit (QFfit) and naïve Bayes approximation for distance adjustments (Supplemental Methods).

### 2.6 WGBS analysis in plasma samples from patients with and without hepatocellular carcinoma

In the comparison of plasma samples from individuals with (n = 26) or without (n = 32) hepatocellular carcinoma (HCC) [33], for each sample, we extracted sequenced reads overlapping the genomic intervals within 200 bp from each of the panels’ CpG sites. We calculated the average methylation of all sequenced CpGs in a region. Thus, we separately added up unmethylated and methylated CpGs across all the individual reads and denoted those counts as <u>c</u>ounts <u>u</u>nmethylated and <u>c</u>ounts <u>m</u>ethylated (cu and cm, respectively). Next, given the cu and cm values, we calculated the signal fraction, x, for each sample at each locus. We calculated x as cm/(cu+cm) when methylation levels in reference plasma samples were close to 0 or as cu/(cu+cm) when methylation levels in reference plasma samples were close to 1. Since tumor signal in plasma is diluted, binarization thresholds at each target locus were set as the highest observed x-value in the 32 control plasma samples. If a given plasma sample from a patient with HCC had an x-value above the threshold, it was assigned a value of 1, otherwise it was assigned a value of zero.

### 2.7 Plasma collection from donors with and without cancer

Plasma samples (up to 4 mL) from donors with colon cancer (n = 20), pancreatic cancer (n = 24), liver cancer (n = 26), prostate cancer (n = 23), stomach cancer (n = 21), and healthy individuals (n = 32) were purchased from Fox Chase Cancer Center (Philadelphia, PA) and Audubon Bioscience (Houston, TX), or were acquired from patients at the NIH Clinical Center (prostate cancer, n = 7) after approval by the Institutional Review Board (IRB) (IRB#16-C-0010/1ZIABC011680) as part of the clinical trial registered as NCT02594202. All patients provided written informed consent in accordance with the principles of the Declaration of Helsinki. In brief, whole blood was collected in EDTA, ACD, or Streck tubes and processed for plasma isolation within 4 hours. The plasma isolation protocol included double centrifugation at 15,000xg for 15 min. at 4°C. Plasma was stored at - 80°C until DNA extraction. These samples were tested using the 8 and 39-marker panels.

### 2.8 Extraction of cfDNA from plasma samples

DNA was extracted from plasma samples using either the NeoGeneStar^TM^ (Somerset, NJ) Cell Free DNA Purification Kit or the Qiagen (Hilden, Germany) QIAamp® Circulating Nucleic Acid kit (Qiagen, cat# 55114). For NeoGeneStar^TM^ processed samples, saline solution was added to the plasma samples to reach a total volume of 5mL as input before proceeding according to the manufacturer’s instructions. DNA was eluted in 23 µl DNase-free water. For Qiagen extracted samples, we followed a modified version the manufacturer’s instruction, including mixing of 4mL plasma with 400µl Proteinase K and 3.2 mL Buffer ACL (containing 1µg carrier RNA) and incubating this at 60°C for 30 min. After adding 7.2 mL Buffer ACB and incubating on ice for 5 min., the samples were passed through the QIAvac 24 Plus manifold. Upon a single wash step using the vacuum system, we transferred all columns to collection tubes and performed the remaining part of the protocol using centrifugation (15,000 – 20,000 x g). cfDNA was eluted in 30µl DNase-free water in a two-step elution by re-applying the eluted material to the column. The total cfDNA output across samples can be viewed in Table S1.

### 2.9 Library preparation and probe hybridization for targeted methylation sequencing

We utilized the Twist NGS Methylation Detection System (Twist Bioscience, San Francisco, CA) for probe design and target enrichment of the 8 and 39-marker detection panels. Initially, cfDNA libraries were prepared according to the NEBNext Enzymatic Methyl-seq Library Preparation Protocol (Twist Bioscience). In brief, DNA was end-prepped and ligated to universal EM-seq adapters followed by a 2-step enzymatic conversion of non-methylated cytosines to uracil utilizing the TET2 and APOBEC enzymes. Converted DNA was PCR-amplified by 10 cycles using unique barcode primers for each sample. Next, the Twist Targeted Methylation Sequencing Protocol was applied for probe hybridization capture and enrichment according to the Twist Bioscience protocol. In brief, libraries were pooled and mixed with the custom methylation panel containing 120 bp biotinylated probes targeting the relevant CpG sites in each of the biomarker panels. To capture the widest range of methylation differences, two capture probes were designed for each locus, each intended to hybridize with either fully methylated or fully unmethylated molecules. Additionally, the experimental design allowed for capture of partially methylated DNA molecules. The probe hybridization reaction went for 2-4 hours.

### 2.10 Targeted methylation sequencing and analysis of plasma samples

Enriched DNA libraries were sequenced using either the NextSeq 2000 (200 cycles) or NovaSeqX (300 cycles) platform (Illumina). FASTQ files were subjected to FastQC and reads with a mean quality score < 20 were discarded. Adapter sequences were trimmed using Cutadapt version 3.7 and reads were aligned to the genome (hg19) using Bismark version 0.22.3. Methylation status was assigned using Bismark’s methylation extractor with parameters “--paired-end --no_overlap --ignore 4 --ignore_r2 3 --ignore_3prime_r2 1”. To assess the efficiency of conversion of unmethylated cytosines to uracil (ending up as thymine after PCR amplification) we evaluated the degree of methylation at non-CpG sites based on all reads regardless of target (only excluding reads mapping to the mitochondrial DNA). If methylation across all non-CpG sites was > 3%, the sample was discarded. As an additional safeguard, we filtered out reads in which more than 8 non-CpG cytosines were reported as methylated (e.g., not converted). Finally, we filtered out reads spanning < 90% of CpGs expected to be present in the ROI. A list of the 8 and 39 regions and the number of CpGs contained within each region can be found in Table S2. Percent methylation was calculated as the number of methylated CpGs divided by the total number of CpGs in that read. If a read covered < 100% of CpGs in a given region we made the assumption that missing CpGs were unmethylated to have a consistent denominator. To test the performance of both biomarker panels, we applied a weighted ensemble machine learning model. The feature input was constructed as cumulative percentage of reads at or below each methylation-density bin for a region defined within each probe. To ensure that minority-class observations were maximally represented in the training data for each split, we used leave-one-out (LOO) cross-validation. For the 8-marker panel training data, we applied SMOTEENN [38] to synthetically augment the minority class and to remove putative noisy samples. Models were trained with the AutoGluon-Tabular [39] framework using the best quality preset and a 5-minute training budget per LOO fold; extending the budget to as long as 1 hour per fold did not meaningfully improve validation performance. For the 8-marker panel, we defined a custom evaluation metric that prioritizes specificity until the number of misclassified non-cancer samples in the validation set exceeds 1. After this threshold, the metric transitions to a heavily sensitivity-weighted combination of sensitivity and specificity. For the 39-marker panel, balanced accuracy was used for model evaluation. All reported performance metrics were computed from predictions on the held-out test set.

## 3 Results

### 3.1 Performance of the DNA methylation marker panels in detecting tumors and classifying tumor type

The main goal of our study was to identify a limited number of methylation sites that can be used to (1) detect the presence of cancer in a sample, and (2) distinguish among the 13 major tumor types from the TCGA repository of 4,052 samples (Table 1). Colon and rectal cancer samples were previously reported as being nearly indistinguishable [30], therefore we combined them into one group, CRAD, for our analyses. Methylation array data from an additional 2,711 peripheral blood reference samples (Table 1) were used to establish background methylation levels expected in blood draws from healthy individuals. Using these data, we initially identified a set of 8 loci whose methylation profiles not only distinguished tumor samples from healthy blood samples but also exhibited similarities between healthy tissues and blood samples from cancer-free individuals (see Methods). This requirement establishes that methylation at these loci is altered in tumors and does not appear in the background signal generated from healthy blood. Thus, in the context of binary tumor classification, the methylation patterns deviate from what is observed in normal conditions. When we applied this 8-probe panel to methylation data from individual tumors of each cancer type in our TCGA discovery dataset, it identified 91% of the tumor samples (Figure 1A). The panel detected > 95% of samples for four specific tumor types: colon and rectal (combined), stomach, and uterine tumors, whereas the detection rate was lowest for pancreatic tumors, at 74% (48/65). Methylation data from all peripheral blood samples of individuals without cancer were correctly identified as non-tumor samples, in alignment with these samples being used as a reference (Figure 1A). Similarly, 99% of normal tissue samples were accurately identified as non-tumor samples (Figure 1B). The false-positive rate was zero for most normal tissue samples, including stomach, pancreas, lung (squamous cell and adenocarcinoma), kidney (clear cell and papillary cell), liver, and uterus. The highest false-positive rate (6%) occurred in normal prostate tissue. The sample size for some normal tissue types was limited, potentially impacting the accuracy of our false-positive rate estimates.

**Figure 1.**
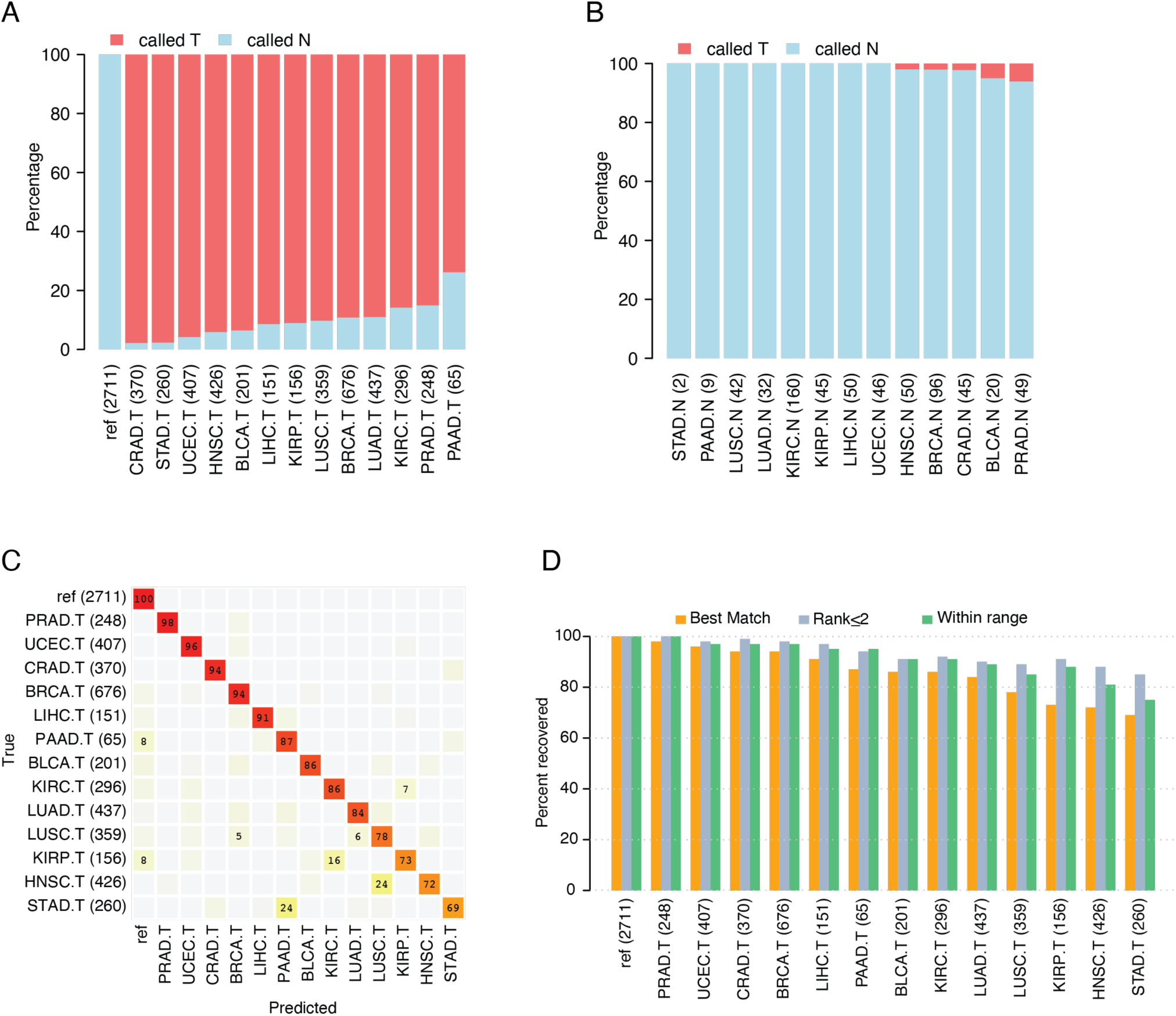
Performance of the 8- and 39-methylation marker panel in the discovery cohort. (A) Performance evaluation using the 8-marker panel on the TCGA tumor dataset, which includes data for 13 tumor types (denoted as “.T”), alongside a reference dataset comprising data from peripheral blood samples ("ref," first column). The percentage of samples identified as tumors shown in red, and those identified as normal samples are shown in blue. (B) Same as in (A) but evaluating TCGA data from 13 normal tissue types (denoted as “.N”). (C) Evaluation performance of the 39-marker tissue classification panel on TCGA tumor dataset, which includes data for 13 tumor types (“.T”) and a reference dataset comprising peripheral blood samples ("ref"). Samples are presented in rows and classification categories in columns. The percentages in each row sum to 100%, with only values ≥5% displayed. (D) Percent recovery of the correct tumor type using three different classification criteria for the 39-marker classification panel, using the same datasets as in (C). TCGA tumor types and sample sizes are labeled as indicated in Table 1. Sample numbers are given in parentheses.

In addition to these 8 loci utilized for tumor-normal calling, we identified 39 CpG sites that could be used for tumor-type classification (see Methods). When applied to the full TCGA collection as a discovery dataset, our 39-marker classification panel returned a median of 86% correct classifications across all 13 tumor types (range 69% - 98%) (Figure 1C). Six tumor types – prostate, uterine, colon and rectal combined, breast, and liver – were classified correctly for more than 90% of samples. Four additional tumor types – pancreas, bladder, kidney clear cell, and lung adenocarcinoma – were classified correctly for 84% - 87% of the samples. Meanwhile, the four remaining tumor types – lung squamous cell, kidney renal papillary cell, head and neck, and stomach – achieved correct classification rates of 69 - 78%. In most instances, the misclassified samples were predicted to be another tumor type originating from the same organ. For example, 6% of lung adenocarcinoma samples were inconsistent with the TCGA labels, being classified as lung squamous cell samples, and 16% of kidney renal clear cell tumor samples were inconsistently classified as kidney renal papillary cell carcinoma samples (Figure 1C). Interestingly, 24% of head and neck tumors were also incorrectly classified as lung squamous cell tumors. Nevertheless, this classification is plausible, as previous studies using the same dataset show that lung squamous, head and neck squamous, and a subset of bladder adenocarcinomas coalesce into one subtype by gene expression and mutation studies [40, 41]. Finally, 99.9% of the 2,711 peripheral blood reference samples were correctly classified, with only two samples being misclassified (Figure 1C). These findings indicate that the 39-marker panel robustly distinguishes among the 13 tumor types when used on DNA methylation data obtained directly from tumors.

Equipped with knowledge of the site of tumor origin, we experimented with two additional criteria to increase the percentage of tumor samples that were correctly classified (Figure 1D, Figure S2 and Supplemental Methods). The original approach described above utilized a "best match" criterion, where the highest scoring tumor type was selected after comparing a sample to every tumor-type category and the blood reference category. An alternative criterion, called “rank :<2,” encompassed all tumor types that were ranked as either the first- or second-best match for each sample. Using this approach the fraction of correctly classified samples increased from the former median of 86% to 93% (with a range across cancer types spanning from 85%-100%) (Figure 1D and Figure S2). Considering a second alternative criterion, named “within range,” we considered a prediction to be a true positive if it fell within a specified distance range of the sample. For every sample a vector of distances to each tumor class was calculated, and if the tumor type matching the sample fell within the defined range, it would be counted as a correct prediction, irrespective of its’ rank. With this approach the median percentage of correctly identified tumors from the discovery set was 91%, with a range across cancer types from 75%-100% (Figure 1D, Figure S2 and Supplemental Methods). We assessed the distribution of ranks for individual samples across all tumor types as well as distribution of the number of types that fell within the specified range (Figure S3). All of the tumor type categories showed a mean rank < 2, and the average number of tumor types within range was also < 2. Finally, we also we examined the performance of the 39-marker panel on the TCGA discovery dataset using different approaches for measuring distance. This included the utilization of QFfit (Quantile fraction fit) and variations of a naïve Bayes classifier (Bayes and Bayesf; see Figure S4). We conclude that the original distance metric showed the best overall classification performance.

It is important to highlight that the 39-marker panel is specifically designed to predict tumor type, assuming that the tumor is already known to be present. As expected, this panel also demonstrates some capability to classify normal tissue samples by organ (Figure S5), owing to the fact that our tumor classifications are based on their distinctions from blood, rather than their deviations from normal tissues, i.e., the signature is specific to the tissue type, not the tumor presence. Therefore, we recommend employing the 8-marker panel to ascertain whether the sample is tumor or normal in status. Finally, while using the same collection of data for discovery and testing datasets carries a risk of overfitting, we have taken measures to ensure that our analysis remains robust against overfitting, such as assessing training and testing sets (see Supplemental Methods).

### 3.2 Performance of panels when applied to independent validation data

For independent validation, we applied the 8- and 39-marker panels individually to a diverse collection of eight independently generated cancer datasets, which included both methylation array data and WGBS data (Table 2). The collection comprised 908 tumor samples, encompassing various types such as breast, colorectal, kidney clear cell, liver, lung (not specified as squamous cell or adenocarcinoma), pancreas, and prostate tumors. The data were generated from the Illumina 450K methylation array, except for liver tumors, for which WGBS data were available. Additionally, there were data for 275 normal tissue samples from colon, lung, kidney, pancreas, and breast tissue, generated from the Illumina 450K methylation array. Finally, data for 32 plasma samples from individuals without cancer, generated through WGBS, were also included.

The 8-marker panel for distinguishing tumor vs normal phenotype detected 69-93% of the validation tumor samples except for kidney tumors where only 20% were detected (Figure 2A). Notably, data for 3 of the 8 markers were absent in the kidney tumor sample cohort, and two of those missing markers were specifically discriminative for kidney cancers (Figure S6). Excluding kidney tumors, the median detection rate for tumors was 85%. This performance is slightly worse than the rate for our 8-marker discovery set, which was 91%. The 8-marker validation test performed best for prostate and colon tumors, with incorrect classification rates of ≤10%. The panel only misclassified 1 out of 32 WGBS plasma samples from individuals without cancer (i.e., reference samples) as tumor samples (Figure 2A). As few as 10 out of 275 (3.6%) normal validation tissue samples were incorrectly classified as tumor samples (Figure 2B). These misclassifications were limited to pancreas (1 out of 41) and breast (9 out of 159) tissue samples. The majority of incorrectly classified breast tissue samples (8/9) were obtained from tumor-adjacent breast tissue and we therefore cannot rule out the possibility that the tumor-adjacent samples were aberrantly methylated [42].

**Figure 2.**
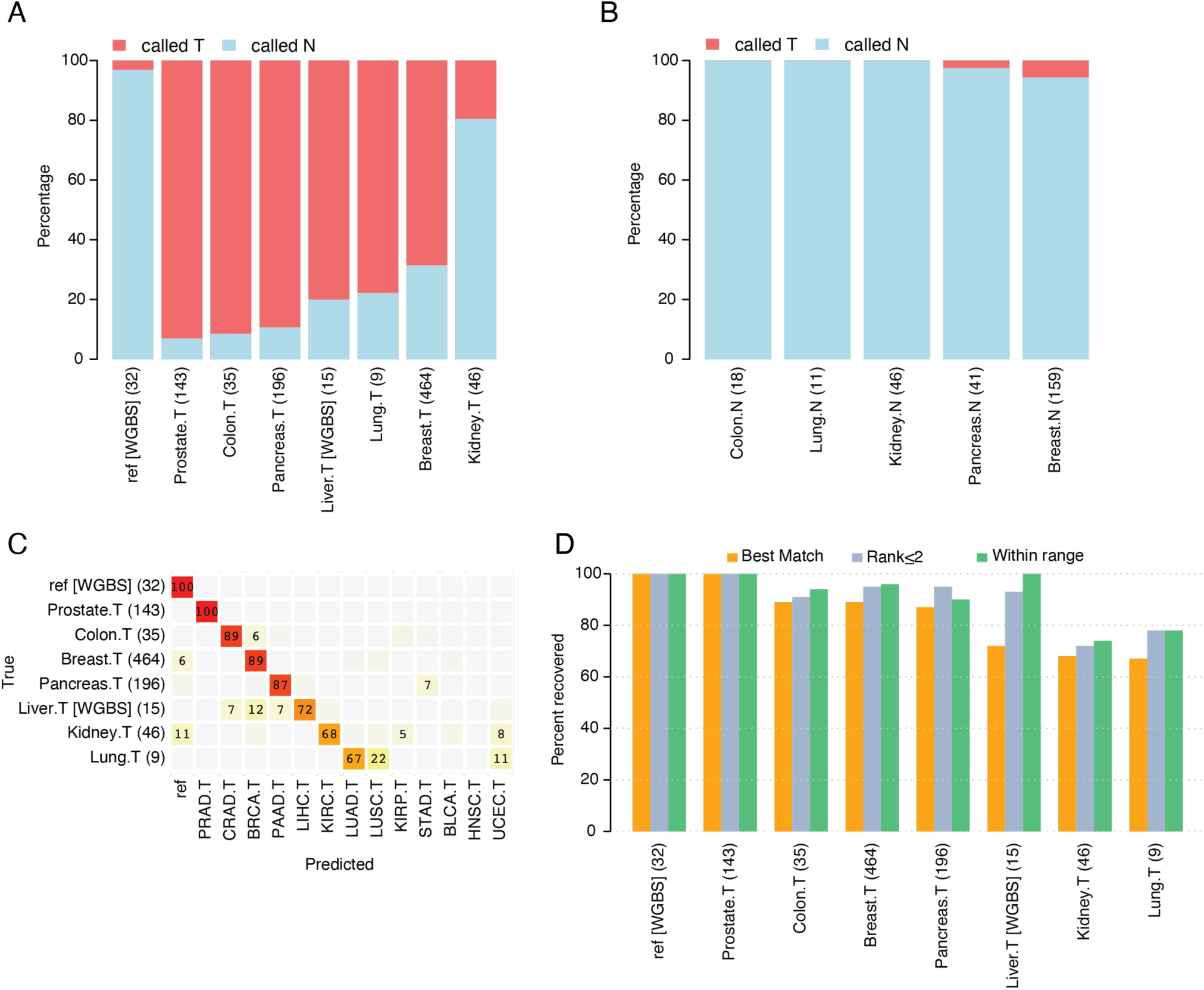
Performance of the 8- and 39-methylation marker panels in independent validation datasets. Both Illumina methylation array data and WGBS data (liver cancer and reference) were considered. (A) Percentage of tumor samples (“.T”) detected correctly by the 8-marker panel; the percentage of samples identified as tumors is shown in red, and as normal samples in blue. The percent of reference samples correctly detected is shown in the leftmost bar. (B) Same as in (A) but evaluating the percentage of normal samples (“.N”) detected correctly by the 8-marker panel. (C) Performance of the 39-marker tissue classification panel shown for validation tumor datasets (denoted “.T”) and a reference set containing healthy blood plasma (denoted "ref [WGBS]"). Samples are presented in rows and classification categories in columns. The percentages in each row sum to 100%, with only values ≥ 5% displayed. (D) Percent recovery of the correct tumor type using three different classification criteria for the 39-marker classification panel, using the same datasets as in (C). The number of samples for each condition is listed in parentheses.

In terms of independent validation of the 39-marker tissue classification panel, performance was comparable to the results achieved with the TCGA discovery dataset. For example, using the “best match” approach, the 39-marker panel correctly classified a median of 87% of tumor validation samples (with a range from 67% to 100%; Figure 2C). This performance is comparable to the discovery-set rate of 86%. Prostate, colon, breast, and pancreas tumor samples were correctly classified between 87% and 100% of the total, mirroring the outcomes observed in the previous results. The validation test’s ability to correctly identify liver, kidney, and lung tumor samples was slightly lower, ranging between 67% and 72%. These rates aligned with the discovery-set rates for kidney and lung tumors; however, classification of liver tumors showed poorer performance in the validation dataset compared to the discovery set. This could likely be attributable to low coverage WGBS data (reported in [33] and [43]). As was true for the discovery dataset, we observed cross-classification of tumor types by organ. For example, 5% of kidney renal clear cell tumor samples were incorrectly classified as kidney renal papillary cell tumors. The actual proportion of lung adenocarcinoma vs. squamous cell tumors was not known; however, 22% of the lung tumor samples (2 of 9) were classified as lung squamous cell (i.e., LUSC). All 32 normal blood plasma samples were correctly classified as reference using the 39-marker panel, consistent with this dataset being used for additional filtering of tissue-specific probes during the selection process (see Supplemental Methods; Figure 2C). We further evaluated the performance of the 39-marker panel by applying the additional classification criteria approaches, “rank ≤2” and “within range” (Figure 2D). Using these criteria improved the correct classification rate for all validation set tumor types for which the “best match” rate was < 100%. For most tumor types, the average number of tumor types recovered using the “within range” criteria was < 1.5 (Figure S3D).

To extend our validation further we also tested the 8- and 39-methylation marker panels in normal and tumor tissue samples derived from eight different tissue sites as well as 13 whole blood samples from healthy individuals using an amplicon bisulfite sequencing approach employing the Fluidigm Access Array (see Supplemental Methods). By calculating the average percent methylation across all CpG sites in each amplicon region and using the same cutoff values and binarization approach as described in the methods section, all 13 healthy blood samples were correctly classified by both the 8 and 39-marker panel. Except from lung and kidney tumor samples, the 39-marker panel correctly identified tissue of origin in 62 – 100% of tumor samples across different tissue types, and the 8-marker panel successfully classified 37/39 tumor samples (see Figure S7). Together this supports the applicability of the two biomarker panels when using various experimental platforms targeting either single or multiple CpG sites seen with Illumina methylation array and amplicon seq, respectively.

### 3.3 Performance of panels on tissue samples from individuals with non-cancer conditions

For optimal performance as screening biomarkers, methylation levels across the 8-marker panel should not be elevated in non-cancer health conditions, were they to leak tissue DNA into the blood stream. To ascertain the effect of such non-cancer conditions on tumor detection performance, we employed the panel to analyze methylation data extracted from tissue samples representing various non-cancer conditions. This dataset consisted of 200 tissue samples from affected individuals and 165 controls (Table 3), encompassing conditions such as type 2 diabetes, asthma, chronic kidney disease, non-alcoholic fatty liver disease, and endometriosis. Except for non-alcoholic fatty liver disease, the datasets also included non-disease tissue samples from the same organ as controls for all conditions (i.e., unaffected). The 8-marker panel classified all samples correctly, as non-tumor samples. This data also included correct calls for peripheral blood samples available for two conditions, Crohn’s disease and ulcerative colitis (GSE32148), further supporting the robustness of the T-N call probes (Table 3).

### 3.4 Assessment of the 8- and 39-marker panels’ performance using WGBS data from plasma cell-free DNA

Considering the panels’ robust performance in classifying tumor tissue, normal tissue, and normal peripheral blood samples, we explored their application in available WGBS data from cfDNA, isolated directly from patient plasma. WGBS data for plasma samples drawn from 26 patients with hepatocellular carcinoma (HCC) (i.e., primary tumors of the liver) were obtained from the study by Chan et al. [33], conducted at Chinese University of Hong Kong. WGBS data were also available for plasma from donors of 32 non-HCC samples, along with 15 solid HCC tumors, used for comparison (also used in Figure 2). The solid HCC tumors demonstrate the signal strength when tumor tissue is sequenced compared to plasma samples.

For each sample, we examined the fraction of methylated CpGs in individual sequence reads, focusing on 200-bp intervals surrounding the original CpG probe sites identified by the 8-marker panel. We retain the Illumina probe name to discuss these regions. Six out of the eight markers showed significant differences in the comparison of plasma samples from cases vs controls (p < 0.05) after applying Holm’s correction (Figure 3). Due to the marker selection approach (See Methods) we did not expect all markers to show differential methylation. All 58 samples were considered as a single training cohort for a simplistic performance assessment. A plasma sample was classified as originating from a tumor-bearing individual if the signal from at least one of the eight markers exceeded the maximum signal observed in all healthy control samples. This thresholding approach resulted in 100% specificity, as no false positives were observed among the controls. Among the 26 plasma samples from HCC cases, elevated signals in at least one of the 8 markers were detected in 16 samples, corresponding to a sensitivity of 61.5%. To assess if the 8-marker panel mistakenly detected any cases among the control samples, we reversed the case and control labels for the plasma samples and repeated the analysis. No cases were detected among the 32 controls. This suggests that the HCC signal detected by the 8-marker panel is specific to ctDNA.

**Figure 3.**
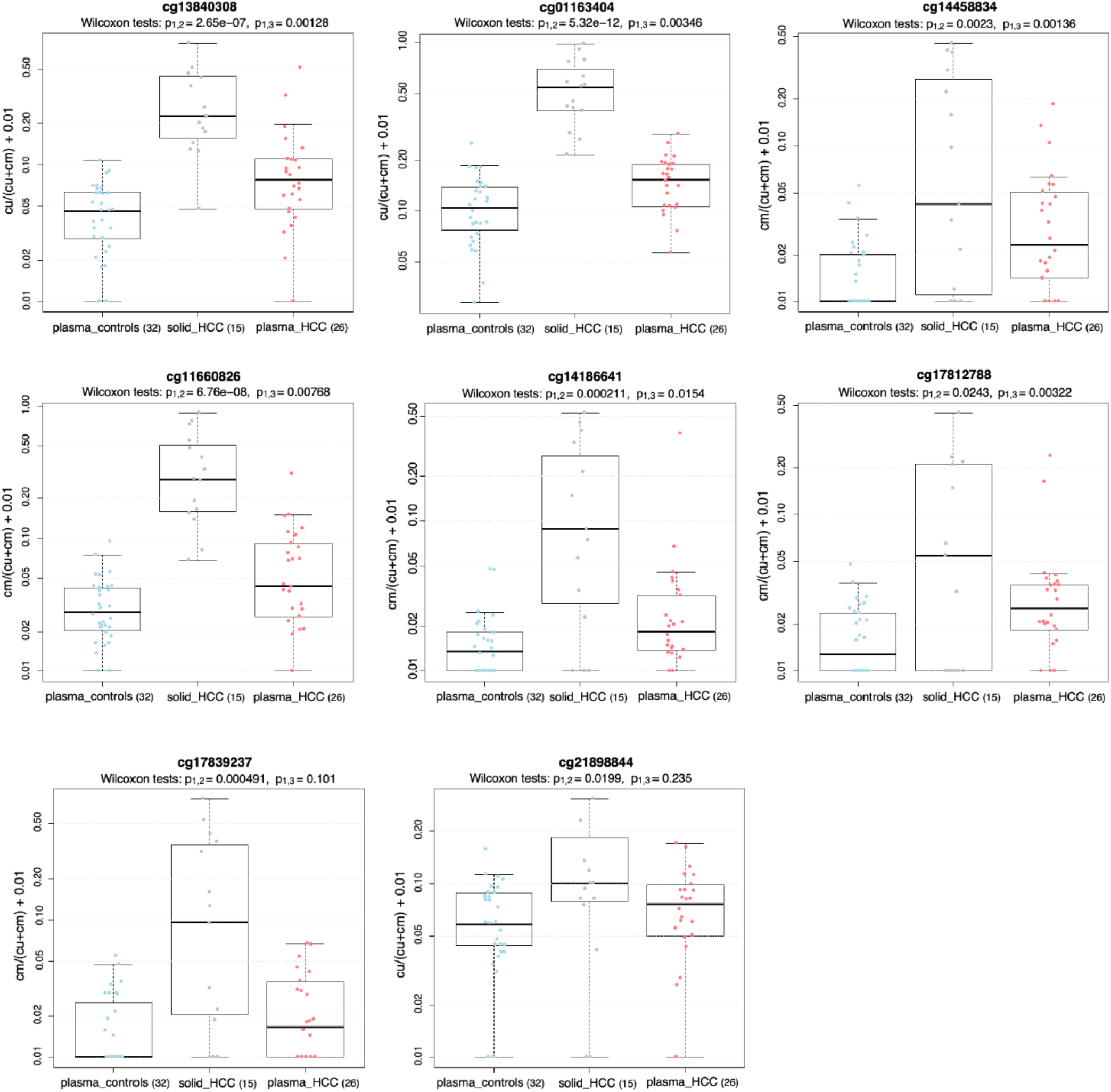
Methylation signal from the 8-marker detection panel using WGBS data from plasma samples of patients with or without hepatocellular carcinoma (HCC). Results from solid HCC samples are provided for comparison. Sample sizes are shown in parentheses. Data reflect the methylation signal from sequence reads within 200-base pairs of each CpG originally annotated as an Illumina probe site. Box plots display the average CpG methylation fraction in DNA from each sample type, with all sequence reads included in the denominator. The difference in methylation signal between plasma samples from HCC patients and those without was calculated using a Wilcoxon rank sum test for statistical significance (p_1,3_).

The performance of the 39-marker panel at classifying these samples by tumor type was less effective on this data. The majority of plasma samples from patients with HCC (25/26) were undetectable as tumor samples and misclassified as references. The explanation stems from the necessity of having concordant signals at multiple tissue-defining loci for robust classification. However, since this WGBS dataset provides shallow coverage, it is insufficient for our purposes. Our previous calculations showed an average read depth of 8-10X in these samples [43]; consistent with the estimate of Chan et al [33]. The low coverage combined with extremely dilute tumor signal in plasma samples likely caused the underperformance of the 39 markers in this data. Indeed, we validated that WGBS data derived from tumor tissue and cell line sources enables classification of sample origin (see Figure S8 and Supplemental Methods).

### 3.5 Performance of panels when applied to independent plasma validation data using targeted methylation sequencing

To accommodate the lack of sequencing depth observed when utilizing WGBS data for analysis of the methylation markers, we implemented a custom methylation hybridization capture panel (Twist Bioscience), enabling enrichment of methylated target sequences in the cfDNA prior to amplification and sequencing. Plasma samples were collected from individuals diagnosed with colon cancer (n = 20), liver cancer (n = 26), pancreatic cancer (n = 24), prostate cancer (n = 23), stomach cancer (n = 21) and those without any cancer diagnosis (n = 32) (outlined in Table 4). Capture was performed on total cfDNA with median input ranging from 35.6 ng (prostate cancer) to 60.3 ng (liver cancer) from 1-4 mL of plasma (Table S1)

**Table 4.** Patient and healthy donor clinical information.

|  | Colon cancer | Liver cancer | Pancreatic cancer | Prostate cancer | Stomach cancer | Healthy controls |
| --- | --- | --- | --- | --- | --- | --- |
| **No. of patients** | 20 | 26 | 24 | 23 | 21 | 32 |
| **Age, median (range)** | 60 (33-82) | 65 (40-74) | 60 (38-78) | 61 (49-75) | 66 (38-80) | 49 (40-68) |
| **Sex, n(%)** |  |  |  |  |  |  |
| Male | 17 (85%) | 10 (58%) | 15 (62.5%) | 23 (100%) | 16 (76%) | 19 (59%) |
| Female | 3 (15%) | 6 (42%) | 9 (37.5%) | - | 5 (24%) | 13 (41%) |
| **Pathologic stage group** |  |  |  |  |  |  |
| I | - | 5 | 1 | - | - | - |
| II | 5 | 4 | 9 | 1 | 8 | - |
| III | 13 | 2 | 1 | 12 | 6 | - |
| IV | 1 | 4 | 12 | 2 | 6 | - |
| NA | 1 | 11 | 1 | 1 | 1 | - |
| **Grade group<sup>a</sup>** |  |  |  |  |  |  |
| 1 | - | - | - | 1 | - | - |
| 2 | - | - | - | 4 | - | - |
| 3 | - | - | - | 1 | - | - |
| 4 | - | - | - | 1 | - | - |

<sup>a</sup> 7 prostate cancer patients did not have prostatectomies performed and thus were not assigned pathological stage information. Instead, we report the biopsy-derived grade group for these patients.

Basic data analysis steps post-sequencing included read filtering and quality control assessment (see Methods). We additionally filtered out any reads spanning less than 90% of the CpG sites in the 120bp regions targeted by each probe. The approach minimized potential variability due to fragment size differences, enabling evaluation of the percentage methylation within each read.

Initially, we investigated whether the 8-marker panel could detect tumor signal and thus separating the cancer samples from healthy plasma samples. To build a multi-cancer detection assay, we applied a weighted ensemble machine learning model in a leave-one-out cross-validation setup. As input, we used distributions of read-level methylation levels, reporting the fraction of reads with a given percent methylation or higher (cumulative proportion). For example, for probe region cg01163404 which spans 4 CpG sites, each sample would have a reported fraction of reads with 1 or more CpG sites methylated (e.g. >0% methylation) (feature 1), fraction of reads with 2 or more CpG sites methylated (>25% methylation) (feature 2), and so on. The idea behind this approach is that tumor signal may derive from a population of fragments with similar, but not identical, degrees of methylation. During each split, a five-fold cross-validation would run inside the training subset of samples before testing the held-out sample. In each fold of the 5-fold cross-validation, we allowed only one normal sample to be incorrectly classified, to maintain a high specificity. Evaluating the prediction results in the testing samples, the 8-marker panel obtained a specificity of 81.2% (Figure 4A). Considering the plasma samples from cancer patients, the panel detected 60% as positive for cancer. While the model was trained with all cancer patient samples labelled as ‘tumor’, we evaluated detection efficiency within each cancer type after completing predictions. The assay detected > 50% of samples in all cancer types, except prostate cancer (17.4%) (Figure 4A). Highest detection rate was observed within stomach cancer (85.7%). To assess the ability of the 8-marker panel to detect early-stage cancer, we extracted prediction outcomes for patient samples categorized as stage I or II (excluding prostate cancer). Out of 32 early-stage patients, the assay successfully detected 25 (78.1% sensitivity) suggesting an ability to capture tumor signal independent of stage. Of note, our liver cancer cohort contained 6 cholangiocarcinoma patients (Table S4), a rare form of cancer with only 3% 5-year survival rate if detected after metastatic spread. Our assay correctly classified 5 out of these 6 cases, including two early-stage patients. Out of the 23 prostate cancer patients included in this study, 7 were enrolled in active surveillance and thus were not yet diagnosed with clinically significant prostate cancer requiring surgery (see Table 4). Interestingly, our 8-marker panel successfully detected 4/7 (57.1%) of these. Overall, the 8-marker panel enabled binary classification of cancer with an AUC of 0.783 (Figure 4B).

**Figure 4.**
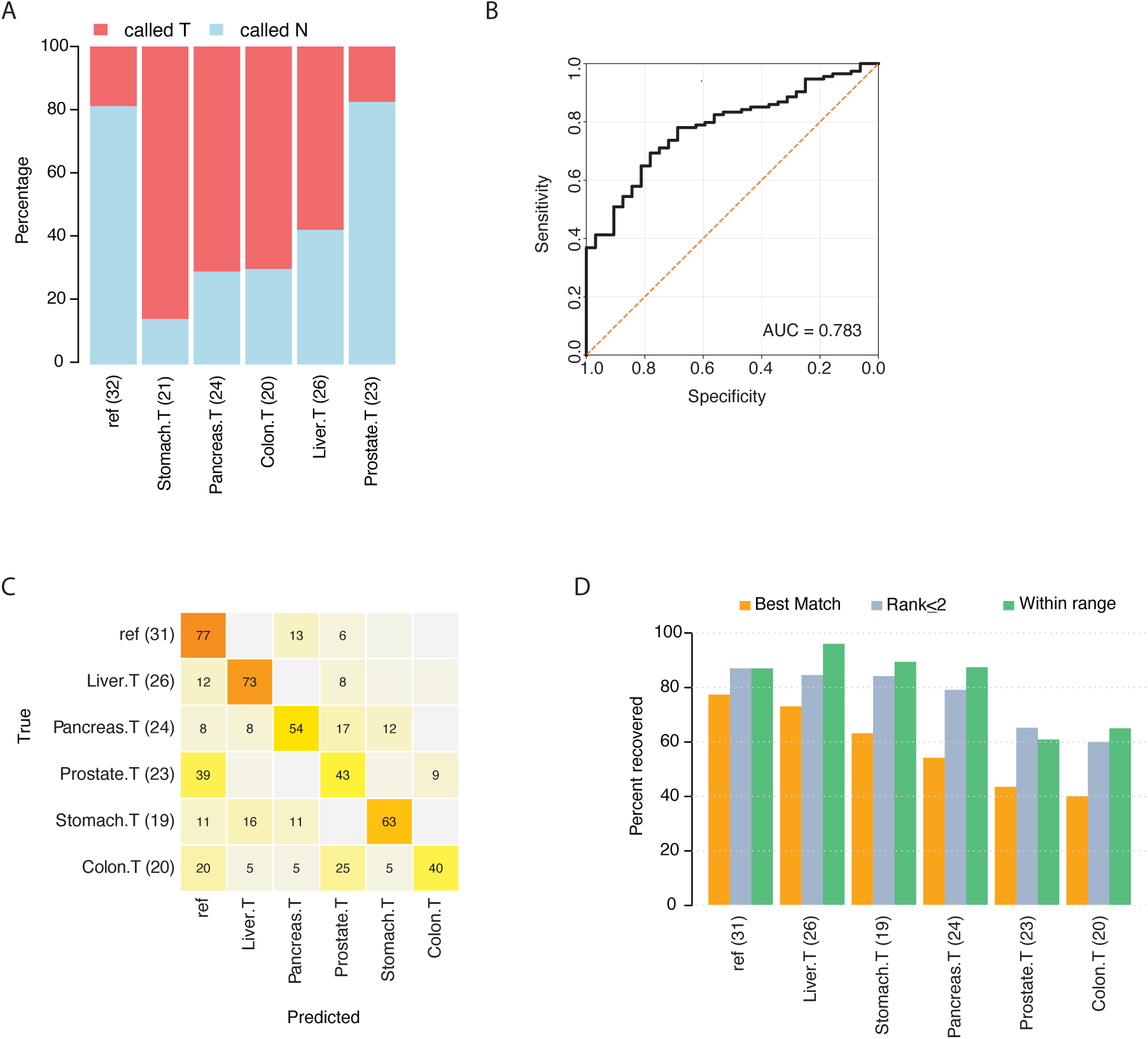
Performance of 8- and 39-methylation marker panels in plasma cfDNA samples using targeted methylation sequencing. (A) Prediction performance of the 8-marker panel in a leave-one-out cross-validated weighted ensemble machine learning model. Plasma samples were derived from patients with 5 different cancer types (denoted as “.T”) alongside reference samples from healthy individuals (“ref”, first column). The percentage of samples identified as tumors shown in red, and those identified as normal samples are shown in blue. (B) Receiver Operating Characteristics (ROC) curve for the 8-marker panel in classifying cancer patient and healthy plasma samples (in total 146 samples). (C) Evaluation performance of the 39-marker tissue classification panel in a leave-one-out cross-validated weighted ensemble machine learning model. Included plasma samples were the same as in (A) except three excluded samples (missing data). Samples are presented in rows and classification categories in columns. The percentages in each row sum to 100%, with only values ≥ 5% displayed. (D) Percent recovery of the correct tumor type using three different classification criteria for the 39-marker classification panel, using the same datasets as in (C). Sample numbers are given in parentheses.

Next, we applied the same type of feature input from the 39 markers in a multiclass machine learning model using balanced accuracy as evaluation metric. Three samples (two stomach cancer and one healthy sample) were excluded due to missing data on some of the probes. In a leave-one-out cross-validation setup, each sample was labelled with risk scores for each category (each of the five tumor types and reference). Using only the best match (maximum risk score) for classification, the panel returned an average of 54% correct classification across all five tumor types (excluding reference samples) (Figure 4C). Performance ranged from 40% for colon cancer samples to 73% for liver cancer samples. A considerable proportion of prostate and colon cancer samples were classified as reference, likely indicating low signal from the tumor tissue of origin in these samples, while mis-classified pancreatic cancer samples had a more even distribution across multiple tissue types. Interestingly, the five colon cancer samples that were incorrectly classified as originating from the prostate were all male donors with age range 58-68, whereas the four samples that classified as reference were from individuals aged 33-53. Thus, it is possible that benign prostatic disease could have contributed with cfDNA from this organ, interfering with the prediction result. Of note, out of the five colon samples classifying as prostate, only one sample was detected by the 8-marker panel. Finally, we tested alternative classification criteria to the best-match approach, allowing the correct tissue type to be ranked ≤ 2 or within range (probability score > 1/6 = 0.17). Best performance was obtained using the ‘within range’ criterion, with an average of 80% correctly classified cancer patient samples with > 85% correct classification of liver, pancreatic, and stomach cancer (Figure 4D). The average number of tumor types retained under the ‘within range’ criterion fell between 1.5-2, while the average rank of correct type varied from 1.5-2.4 (Figure S9), suggesting ‘within range’ as a more efficient diagnostic tool for determining tissue of origin.

## 4 Discussion

Epigenetic biomarkers are useful indicators of tumor types [44–46], subtypes [47–49], presence of disease [50, 51], and responsiveness to therapy [48, 52, 53]. Previously, we showed that *ZNF154* is a multi-cancer biomarker [19, 20] whose differential methylation demonstrates potential clinical utility for tumor diagnostics. In the current study, we focused on finding small sets of markers whose methylation could eventually be used in a liquid biopsy multi-cancer test. Although new markers can be easy to propose, judging the utility of biomarkers for multi-cancer detection and classification, requires critical steps to determine their specificity and capacity to identify the correct tumor type. To be clinically viable, candidate multi-cancer biomarkers must be vetted across a variety of cancer types to document disease specificity. This is especially relevant for non-invasive diagnostics, for which the source tissue is indirectly accessed using blood and often present at exceedingly low concentrations.

In this study, rather than testing performance of a single methylation-site biomarker, we tested a method for picking markers that can discriminate among multiple tumor types and expand to accommodate additional tumor types. To enhance our approach, we examined multiple methods for evaluating the performance of both binary predictions for tumor vs. normal samples and tumor-type classification. In addition to performing *in silico* analyses of data from sample repositories, we performed proof-of-principle experiments in whole tissue and patient plasma samples.

Using data from 13 solid tumor types as well as peripheral blood drawn from individuals without cancer, we identified DNA methylation marker panels that could be used to (1) distinguish between tumor and non-tumor samples and (2) classify tumor samples according to type. Our 8-marker detection panel identified over 90% of solid tumor samples, with a negligible false positive rate for reference blood samples. On average, our 39-marker panel correctly classified 85% of solid tumor samples and nearly 100% of the reference blood samples. Perhaps most importantly, the 8-marker panel demonstrated its utility in assessing cfDNA using custom methylation hybridization capture panels. Although our analysis did not specifically examine the markers’ performance on samples from different tumor stages, our TCGA discovery dataset included early-stage tumors, as detailed in Miller et al. [21] and Funderburk et al. [43]. Furthermore, 25 of the 26 HCC plasma samples that we analyzed in this study, the data for which were provided by Chan et al [33], were Barcelona Clinic Liver Cancer (BCLC) stage A, defined as early stage tumors less than 3 cm. Finally, 32 of the plasma samples tested with targeted methylation sequencing derived from patients with early-stage disease (Stage I or II cancer), of which 25 were labelled as ‘tumor’ (78% sensitivity) by the machine learning model. Therefore, the markers show potential to non-invasively detect tumors at stages where interventions could be curative.

Here we champion the use of DNA methylation rather than the mutational profile of DNA. Single mutation-based tests, although technically accurate, are unable to reliably distinguish a tumor’s tissue of origin due to commonality of somatic mutations across tumor types. In our opinion, mutation-based tests can be optimal for following patient progress in tumors of a known type, whereas epigenetic signatures can be complementary or superior to mutational signatures for multi-cancer detection and classification. Prediction of the affected organ can facilitate timely follow up and localized treatment, including surgery. Moreover, the performance of our 39-marker panel on lung and kidney cancers demonstrates that methylation alterations can distinguish between different tumor subtypes within these organs. It is true that some tumors from different tissues might be more similar to one other than to other tumors from the same tissue (i.e., they display cross-cancer signatures). Indeed, the 39-marker panel misclassified some tumors from both the TCGA dataset and independent sources across organ boundaries, suggesting epigenetic similarities that have not been well characterized. Previous studies of the TCGA dataset have reported that head/neck and lung squamous cell cancers share similarities [40]. Our findings were similar, but also included bladder cancers, consistent with Hoadley et al. [41]. As much progress is being made with methylation-based cancer testing [54], ongoing research on the relationship between genetic aberrations and epigenetic disease signatures might facilitate identification of these complementary signatures for use in specific applications.

Our marker selection strategy was based on analysis of methylation array data. The absence of expansive CpG coverage and the lack of information on the methylation status of adjacent CpGs within the same DNA fragment may have caused us to overlook many useful discriminative loci. However, access to data from hundreds of samples in different tumor categories was an important advantage of using the TCGA methylation array dataset. More extensive CpG coverage and information about the methylation status of CpG sites adjacent to identified markers would further benefit the discovery phase.

We had a greater ability to identify the correct tumor type when we did not demand the existence of abnormal methylation at the markers in tumor samples relative to normal tissue samples from the same organ. This makes sense, as it allows us to utilize probes informative for a tissue-specific signature but not necessarily a tumor-specific signature. In other words, markers producing a signal in a normal tissue can be informative. In fact, one might explicitly *discourage* mutually exclusive tumor-normal tissue differences in probe selection for a multi-cancer test. Nevertheless, to guard against elevated normal tissue-specific signatures in blood plasma, a separate mechanism should be employed to establish the presence of a cancer. In this study, this separate mechanism was an 8-marker multi-cancer set of loci to detect abnormal methylation. We also addressed the issue of potential non-malignant tissue vs malignant tissue contributing to blood, by testing samples associated with multiple non-malignant conditions. Importantly, none of these were detected as tumor samples by our 8-marker panel. However, we have not studied this issue in full, as that would necessitate considering large collections of data derived from blood samples gathered from individuals with multiple nonmalignant underlying conditions. That is beyond the scope of our present analysis.

In assessing the 39-marker panel’s performance, we investigated three classification criteria: (1) the top-ranked cancer type (“best match”), (2) the top two-ranked cancer types (“rank ≤2”), and (3) multiple top-scoring cancer types accepted (“within range”). The two latter criteria provided more information and greater likelihood of predicting the correct cancer type, narrowing the options for clinicians, while avoiding stringent determinations that caused loss of sensitivity. For the majority of solid tumor types, the correct tumor type’s rank averaged between the first and second (i.e., 1-1.5) prediction. Similarly, the average number of tumor types retained under the “within range” criterion fell between 1 and 1.5. This is comparable to the observations among cancer patient plasma samples, where the average number of classification types “within range” was between 1.5-2. Hence the “within range” criterion proved to be more economical than the “rank :< 2” criterion (which is defined to retain 2 tumor types), as the former typically yielded fewer than 2 types for further consideration while delivering comparable performance to the latter. The capability of either of these approaches to improve correct classification could be clinically valuable. This is especially relevant in a screening approach where the correct tumor type is not known *a priori*, such as in blood-based assessments that precede other diagnostic modalities. Because the alternative classification criteria may include more than one tumor-type prediction, they would be most helpful when used in concert with diagnostics available for specific tumor types. We initially applied both panels to WGBS data from plasma samples collected from individuals with and without HCC [33]. Overall, we found these data inadequate for the desired detailed analysis of tumor detection and classification performance. The main issue was shallow depth of coverage and the scarcity of sequenced fragments with multiple CpGs. Despite these constraints, the 8-marker panel detected 61.5% of tumors based on data collected from the plasma of patients with HCC and yielded zero false positives. However, we did not perform cross-validation, and the signal across the 39-marker panel was insufficient for tumor type classification. We conclude that strategies employing deeper sequencing coverage WGBS are needed for this kind of analysis in plasma samples, or alternatively, target enrichment approaches. Importantly, our limited marker set abolishes the need for whole genome sequencing and its attendant cost, greatly focusing the analysis to a small set of regions.

In line with this, we also tested the ability of our 8-marker panel to detect ctDNA from up to 4 mL of plasma using hybridization-based targeted methylation sequencing. The 8-marker, cancer detection panel detected 60% of all cancer patient plasma samples, ranging from 17% to 86% sensitivity for prostate and stomach cancer, respectively (with specificity among healthy donors at 81%). Reasons for the relatively low sensitivity in detecting prostate cancer could be technical, such as the slightly lower DNA input amounts retrieved from this cohort, or biological, such as lower ctDNA fractions from patients with this cancer type [13]. We presented the finding from the weighted ensemble machine learning model in a leave-one-out cross-validated setup. The value of cross validation is the measurement of the models’ predictive capacity when tested on data not used for training, however, a larger analysis will be necessary to set universal cancer detection thresholds and independently validate the results.

Furthermore, the 39-marker panel showed a strong potential for classifying tissue of origin based on the dilute cancer-relevant signal found in patient plasma samples. Using the “within range” approach, the panel correctly classified 80% of the patient samples. While most misclassifications using the best-match approach were directed towards the reference designation, a considerable fraction of pancreatic (17%) and colon (25%) cancer plasma samples misclassified as prostate. Of note, the 39 markers were not selected based on tumor-specificity and thus would likely also capture healthy or benign prostate-derived cfDNA in circulation. Thus, further studies including more thorough diagnostic follow-up would be required to fully interpret the signal from the 39-marker panel.

While these results remain preliminary, they greatly demonstrate the potential of this rather limited set of tissue classification markers.

## 5 Conclusions

In summary, we used the 13-cancer type TCGA dataset to select for multi-cancer markers and examined their performance at identifying and classifying tumor samples. We also showed the use of these markers to detect liver tumors in cfDNA from plasma with low-coverage WGBS data at the 8-marker loci. Our approach faced challenges due to the tiny amounts of tumor DNA found in plasma. To enrich for relevant signal, we introduced a new workflow for cfDNA assessment using hybridization panels in a probe-based methylation capture approach and successfully detected more than half of the cancer patient samples while maintaining a high specificity. The principal challenge in the future will be to improve the accuracy of tumor type classification using the minimal input of cfDNA from plasma samples. Other challenges will include the need to accommodate additional tumor types. Developing a targeted, sensitive analysis method to interrogate DNA methylation at a specific set of genomic loci could advance the ability to diagnose and classify tumors by type without the need for whole genome methylation sequencing in plasma samples. Thus, this area of research certainly warrants further investigation.

## Data accessibility

Raw data are available through the TCGA portal and Gene Expression Omnibus, GEO. Experimental data are available upon request.

## Author contributions

G.M., S.B.C., and L.E. conceptualized the study. G.M., L.E., and Y.C.C. developed the methodology around marker selection and testing in TCGA data. S.B.C., S.A.G., B.D.B, L.J.K., and P.K. established the analytical workflow for analysis of targeted methylation sequencing data from plasma cfDNA. G.M., S.B.C., H.M.P., N.J., S.K.F, S.P., and K.F. validated the methodology in additional datasets and plasma sample cohorts. G.M., P.K., and S.B.C. produced plots for data visualization and interpretation. N.C.W., P.A.P, and A.G.S contributed to patient enrollment, sample collection, and processing. L.E. and J.J.L. provided supervision throughout the study. L.E., S.B.C., and J.L.L. provided funding for the study. G.M., S.B.C., and L.E. drafted the original manuscript. G.M., S.B.C., S.A.G., B.D.B., P.K., K.F., S.P., N.C.W., P.A.P., L.J.K., S.K.F., N.J., Y.C.C., H.M.P., A.G.S., J.L.L., and L.E. performed review and edited the manuscript. All authors have read and agreed to the published version of the manuscript.

## Acknowledgements

We thank Abdel Elkahloun and Bayu Sisay of the NHGRI Microarray and Single Cell Genomics Core as well as the NIH Intramural Sequencing Center for sequencing our methylation capture panel data. Furthermore, we would like to thank Dr. Wenyuan Li (UCLA) and Peiyong Jiang (CUHK) for access to the WGBS datasets used in this study. We thank Kristin Harper of Harper Health & Sciences Communications LLC for help editing the manuscript.

## Funding sources and disclosure of conflicts of interest

This research was funded by Carlsberg Foundation to S.B.C. (grant number CF21-0592), and by the Intramural Program of the National Human Genome Research Institute to L.E. (ZIA HG200323), and the Intramural Program of the National Institute of Environmental Health Sciences to JLL (ZIC ES103371), and the Intramural Program of the National Cancer Institute (1ZIABC011680). This research was supported by the Intramural Research Program of the National Institutes of Health (NIH). The contributions of the NIH author(s) are considered Works of the United States Government. The findings and conclusions presented in this paper are those of the author(s) and do not necessarily reflect the views of the NIH or the U.S. Department of Health and Human Services.

The authors declare that they have no competing interests. L.E. and G.M. are listed as inventors on the USPTO Patent Application US-20220290245-A1.

## Ethics Statement

The study was approved by the IRB of National Institutes of Health, which waived the requirement for informed consent because de-identified archival specimens. Seven prostate samples were associated with National Cancer Institute IRB protocol 16-C-0010, which is registered as NCT02594202 at clinicaltrials.gov.

## List of abbreviations

(ctDNA): circulating tumor DNA
(ctDNA): cell-free DNA
(ctDNA): The Cancer Genome Atlas
(GEO): Gene Expression Omnibus
(CpG): 5’-C-phosphate-G-3’
(ROI): Region Of Interest
(ROC): Receiver-Operating Characteristic curve
(AUC): Area Under the Curve
(T2D): Type-2 Diabetes
(WGBS): Whole Genome Bisulfite Sequencing
(genomic DNA): gDNA
(HCC): Hepatocellular Carcinoma

## Supporting information

Additional supporting information may be found online in the Supporting Information section at the end of the article.

## Supplemental Methods

Marker selection strategy; Alternative classification criteria; Comparison of performance by different classification metrics; Addressing overfitting in TCGA and PBref data; Bisulfite amplicon sequencing; Analysis of WGBS data from cell lines and tissue samples

**Fig. S1.** Overview of the binarization process to call sample types based on methylation at the 39-marker set

**Fig. S2.** Illustration of alternative classification criteria

**Fig. S3.** Distribution of ranks and tumor types within range for the discovery and the independent validation datasets

**Fig. S4.** Performance of different type classification metrics on blood reference and TCGA tumors

**Fig. S5.** Performance of the 39 classification markers in normal tissues

**Fig. S6.** Methylation level across the 8-marker panel in TCGA and blood reference datasets

**Fig. S7.** Performance of the 8- and 39-marker panel in amplicon bisulfite sequencing data from normal and tumor tissue samples

**Fig. S8.** Classification performance in WGBS data

**Fig. S9.** Distribution of ranks and tumor types within range for the plasma validation dataset using machine learning-based classification

**Table S1.** Amount of plasma cfDNA used as input for library preparation

**Table S2.** 8 and 39- marker panel probe regions

**Table S3.** Probe status for the 8-marker panel as defined by median-based cutoff in methylation array data

**Table S4.** Additional clinical information on patient cohorts

## Notes

### Competing Interest Statement

The authors have declared no competing interest.

