## Supplemental Figure for "Discovery and performance of DNA methylation panels for cancer detection and classification in blood"

### Slide 1
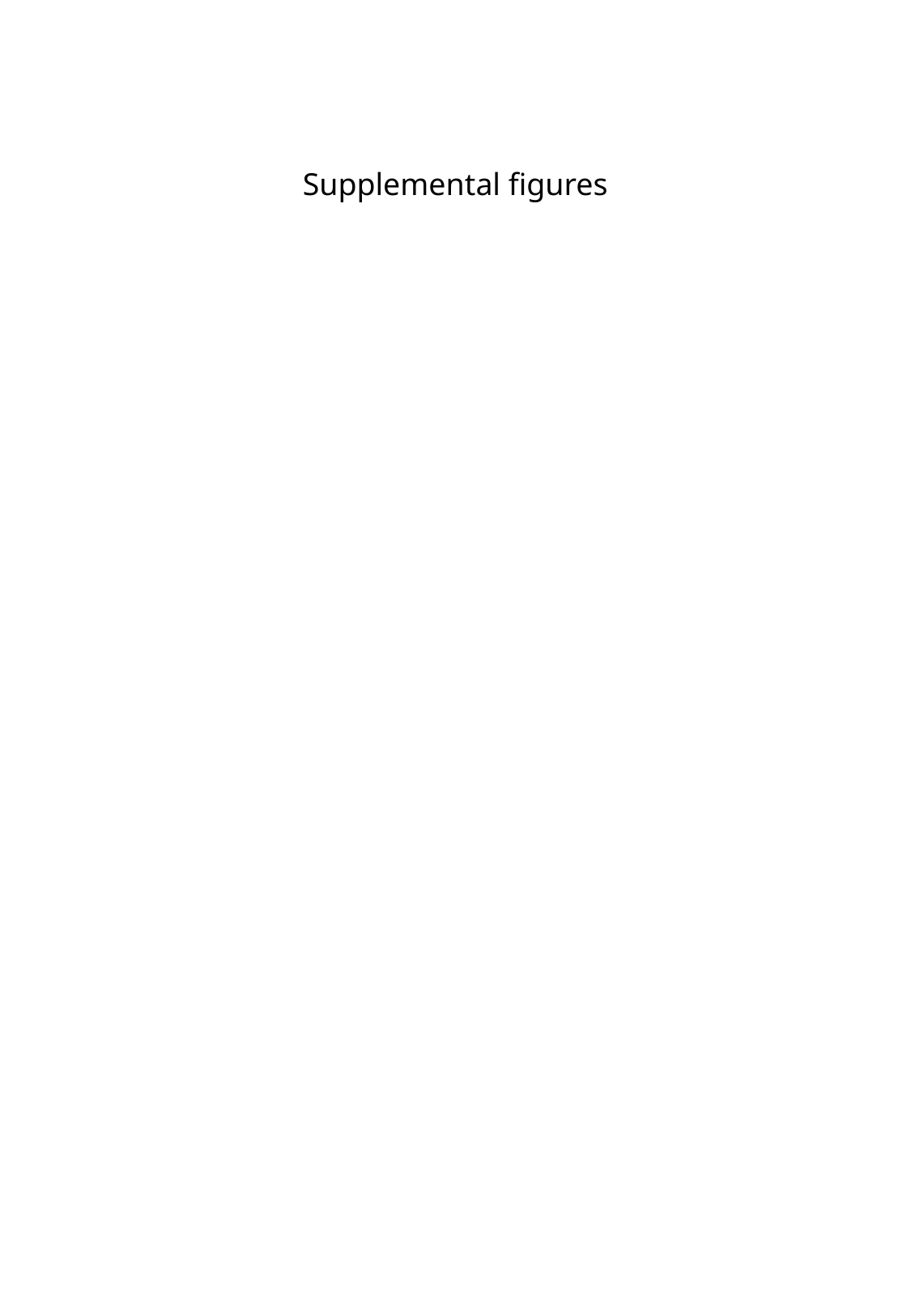

Supplemental figures

### Slide 2
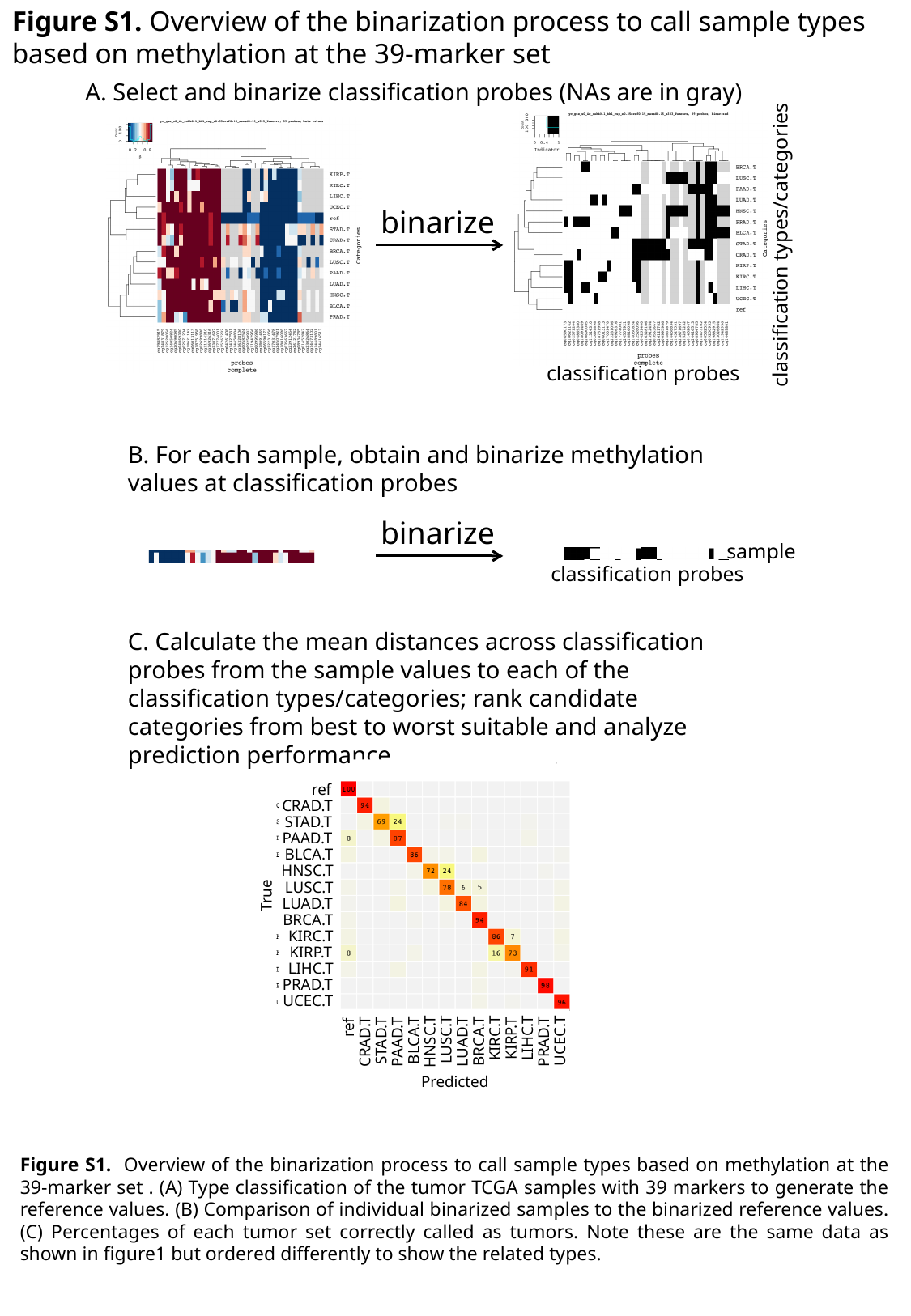

Figure S1. Overview of the binarization process to call sample types based on methylation at the 39-marker set
A. Select and binarize classification probes (NAs are in gray)
binarize
classification types/categories
classification probes
B. For each sample, obtain and binarize methylation values at classification probes
binarize
sample
classification probes
C. Calculate the mean distances across classification probes from the sample values to each of the classification types/categories; rank candidate categories from best to worst suitable and analyze prediction performance
ref
CRAD.T
STAD.T
PAAD.T
BLCA.T
HNSC.T
LUSC.T
LUAD.T
BRCA.T
KIRC.T
KIRP.T
LIHC.T
PRAD.T
UCEC.T
True
ref
CRAD.T
STAD.T
PAAD.T
BLCA.T
HNSC.T
LUSC.T
LUAD.T
BRCA.T
KIRC.T
KIRP.T
LIHC.T
PRAD.T
UCEC.T
Predicted
Figure S1. Overview of the binarization process to call sample types based on methylation at the 39-marker set . (A) Type classification of the tumor TCGA samples with 39 markers to generate the reference values. (B) Comparison of individual binarized samples to the binarized reference values. (C) Percentages of each tumor set correctly called as tumors. Note these are the same data as shown in figure1 but ordered differently to show the related types.

### Slide 3
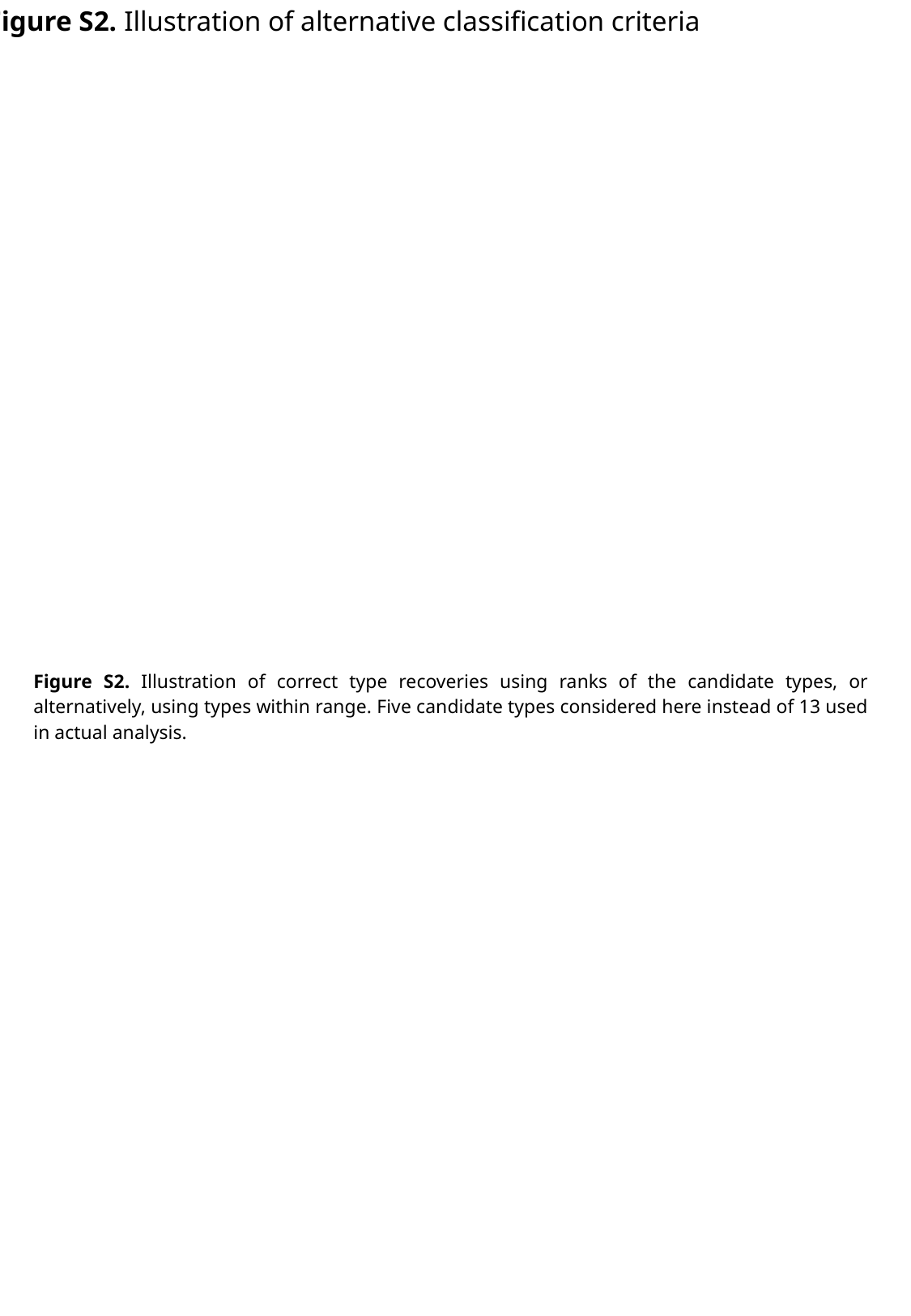

Figure S2. Illustration of alternative classification criteria
Figure S2. Illustration of correct type recoveries using ranks of the candidate types, or alternatively, using types within range. Five candidate types considered here instead of 13 used in actual analysis.

### Slide 4
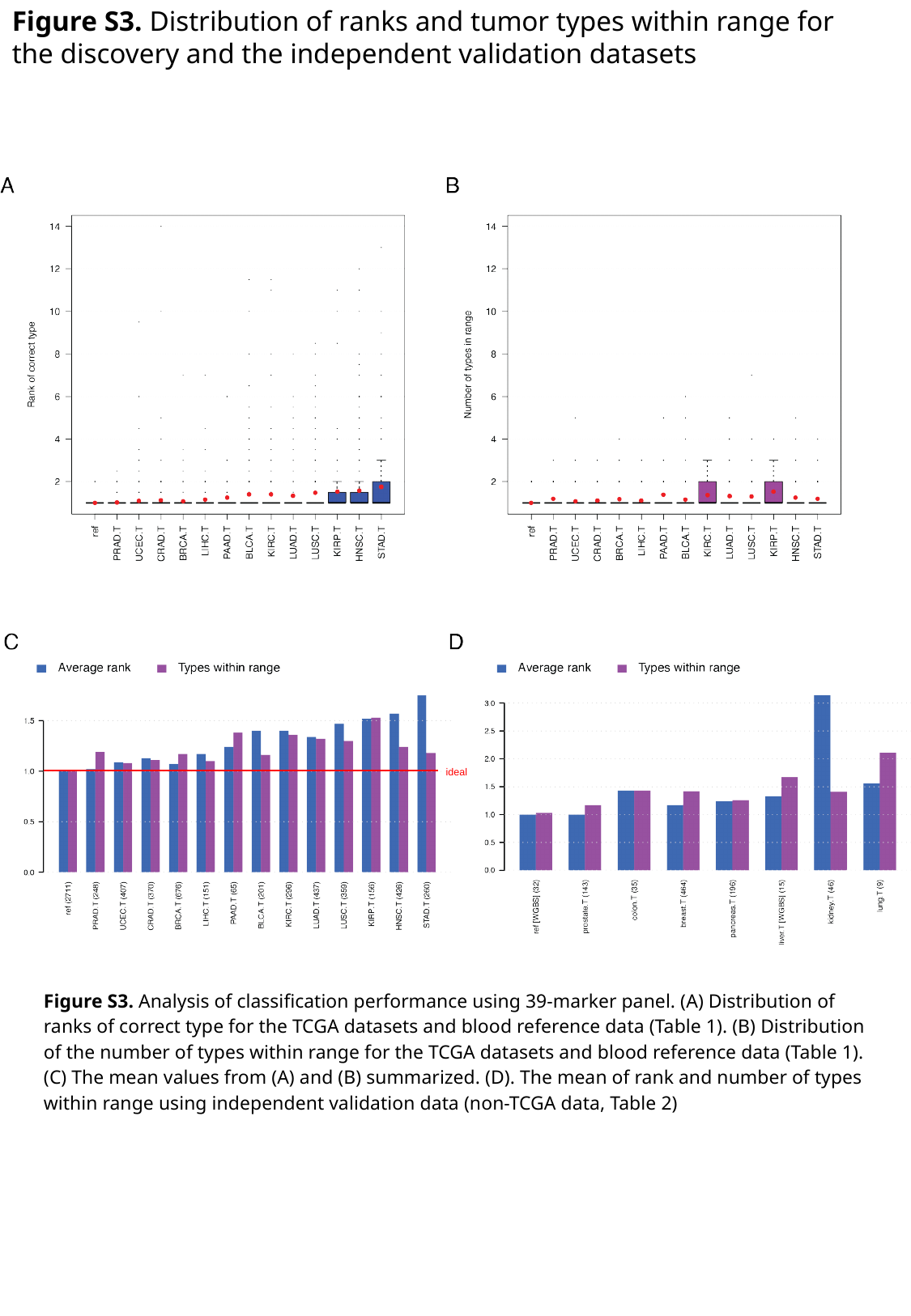

Figure S3. Distribution of ranks and tumor types within range for the discovery and the independent validation datasets
ideal
Figure S3. Analysis of classification performance using 39-marker panel. (A) Distribution of ranks of correct type for the TCGA datasets and blood reference data (Table 1). (B) Distribution of the number of types within range for the TCGA datasets and blood reference data (Table 1). (C) The mean values from (A) and (B) summarized. (D). The mean of rank and number of types within range using independent validation data (non-TCGA data, Table 2)

### Slide 5
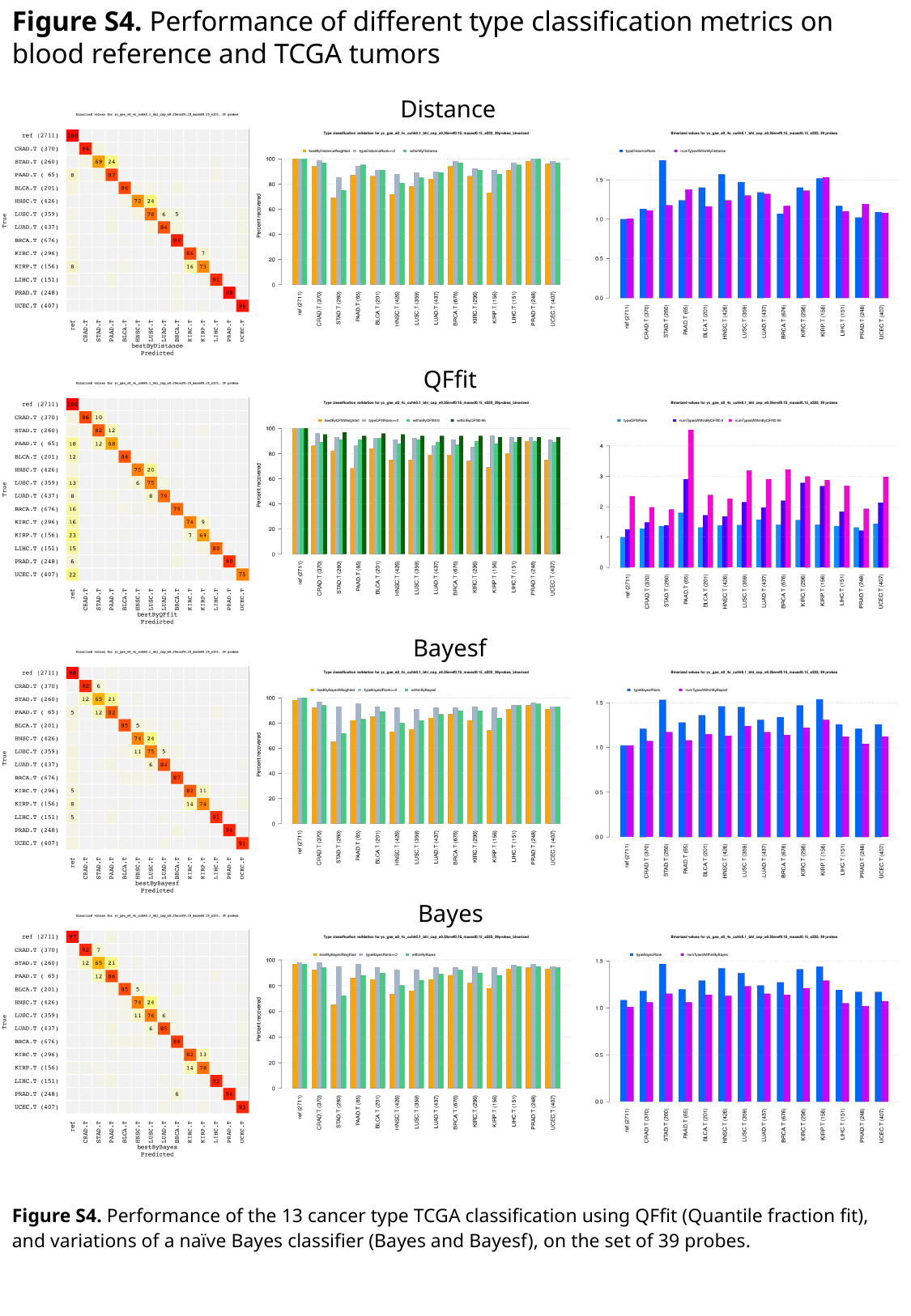

Figure S4. Performance of different type classification metrics on blood reference and TCGA tumors
Distance
QFfit
Bayesf
Bayes
Figure S4. Performance of the 13 cancer type TCGA classification using QFfit (Quantile fraction fit), and variations of a naïve Bayes classifier (Bayes and Bayesf), on the set of 39 probes.

### Slide 6
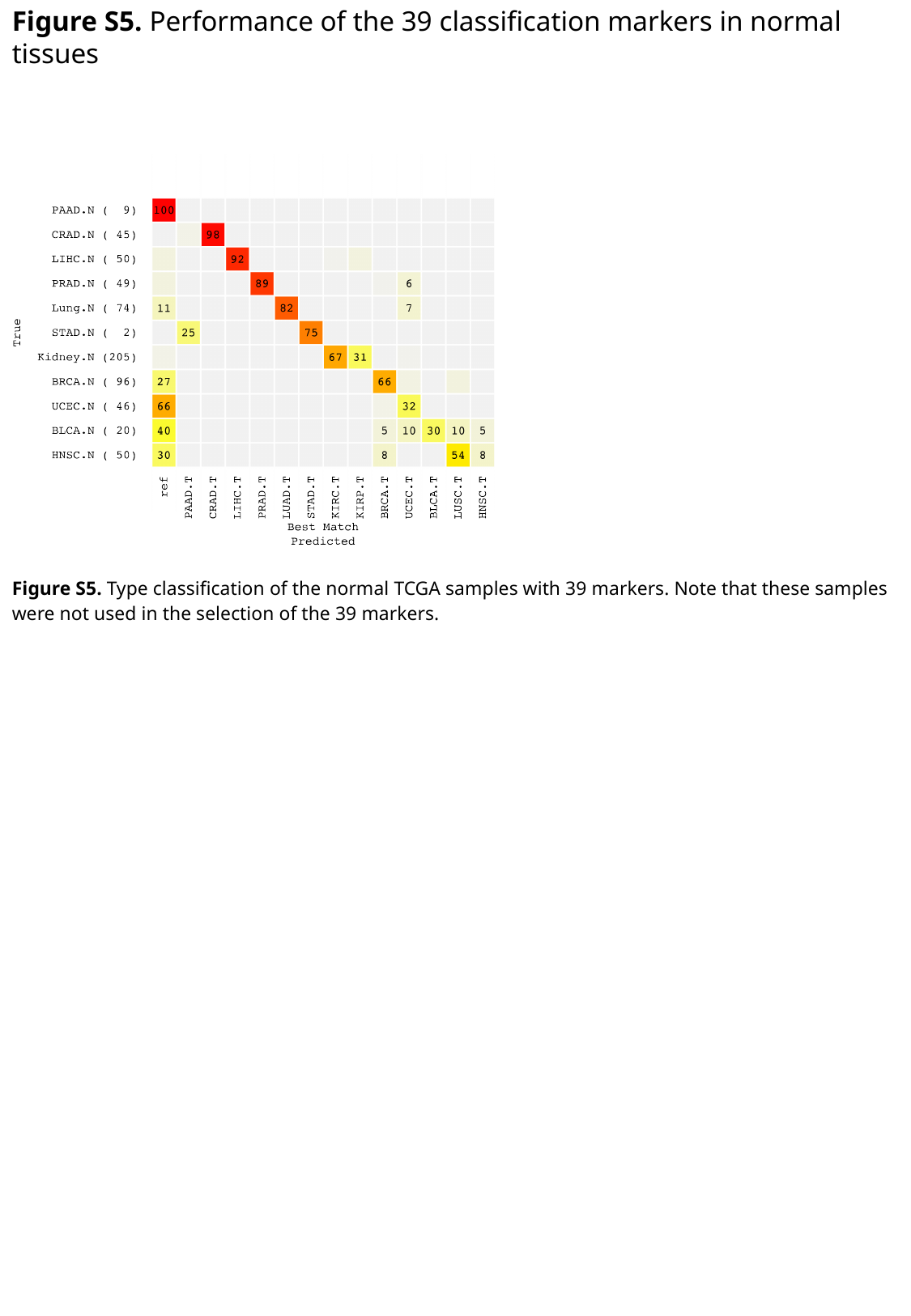

Figure S5. Performance of the 39 classification markers in normal tissues
Figure S5. Type classification of the normal TCGA samples with 39 markers. Note that these samples were not used in the selection of the 39 markers.

### Slide 7
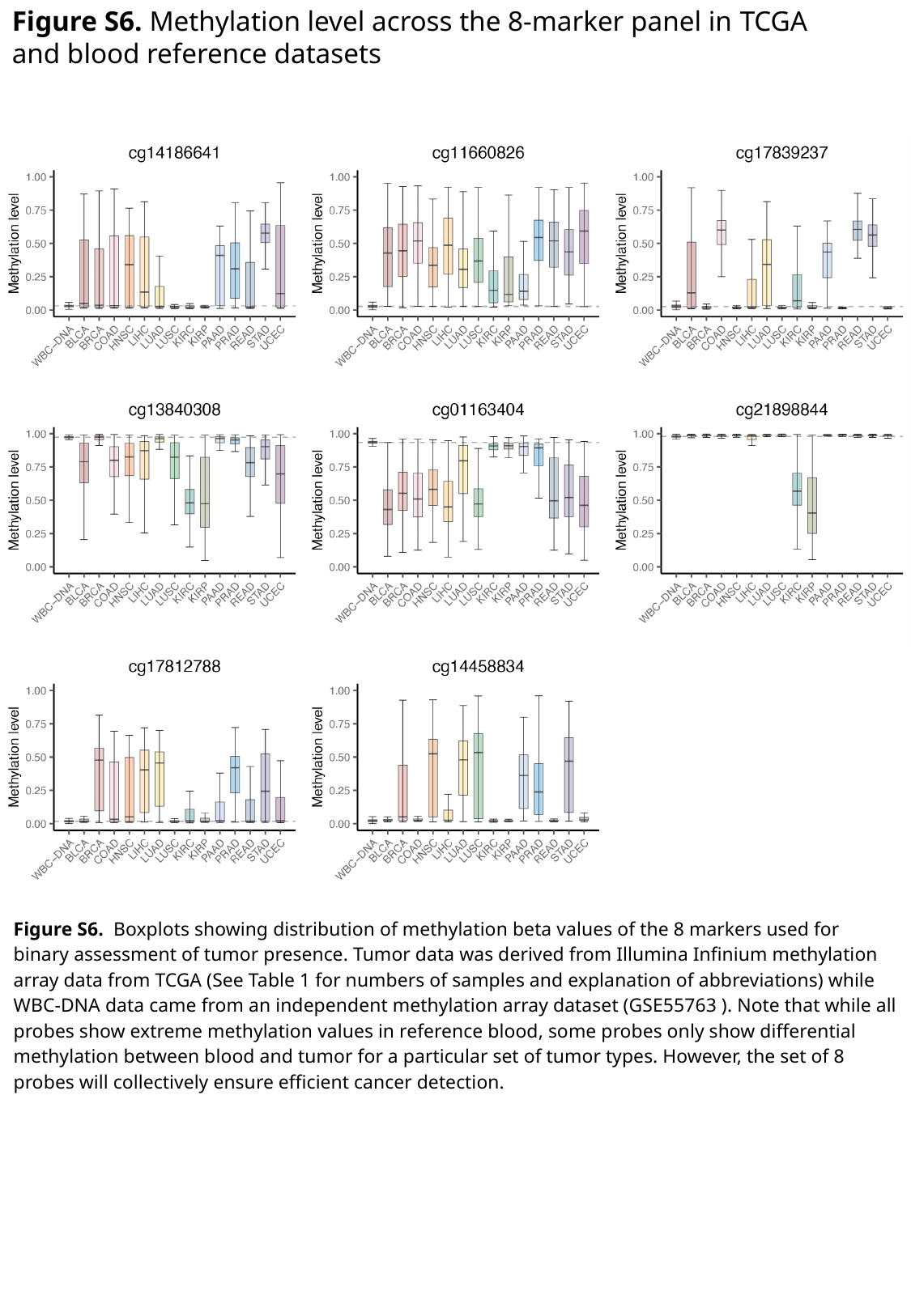

Figure S6. Methylation level across the 8-marker panel in TCGA and blood reference datasets
Figure S6. Boxplots showing distribution of methylation beta values of the 8 markers used for binary assessment of tumor presence. Tumor data was derived from Illumina Infinium methylation array data from TCGA (See Table 1 for numbers of samples and explanation of abbreviations) while WBC-DNA data came from an independent methylation array dataset (GSE55763 ). Note that while all probes show extreme methylation values in reference blood, some probes only show differential methylation between blood and tumor for a particular set of tumor types. However, the set of 8 probes will collectively ensure efficient cancer detection.

### Slide 8
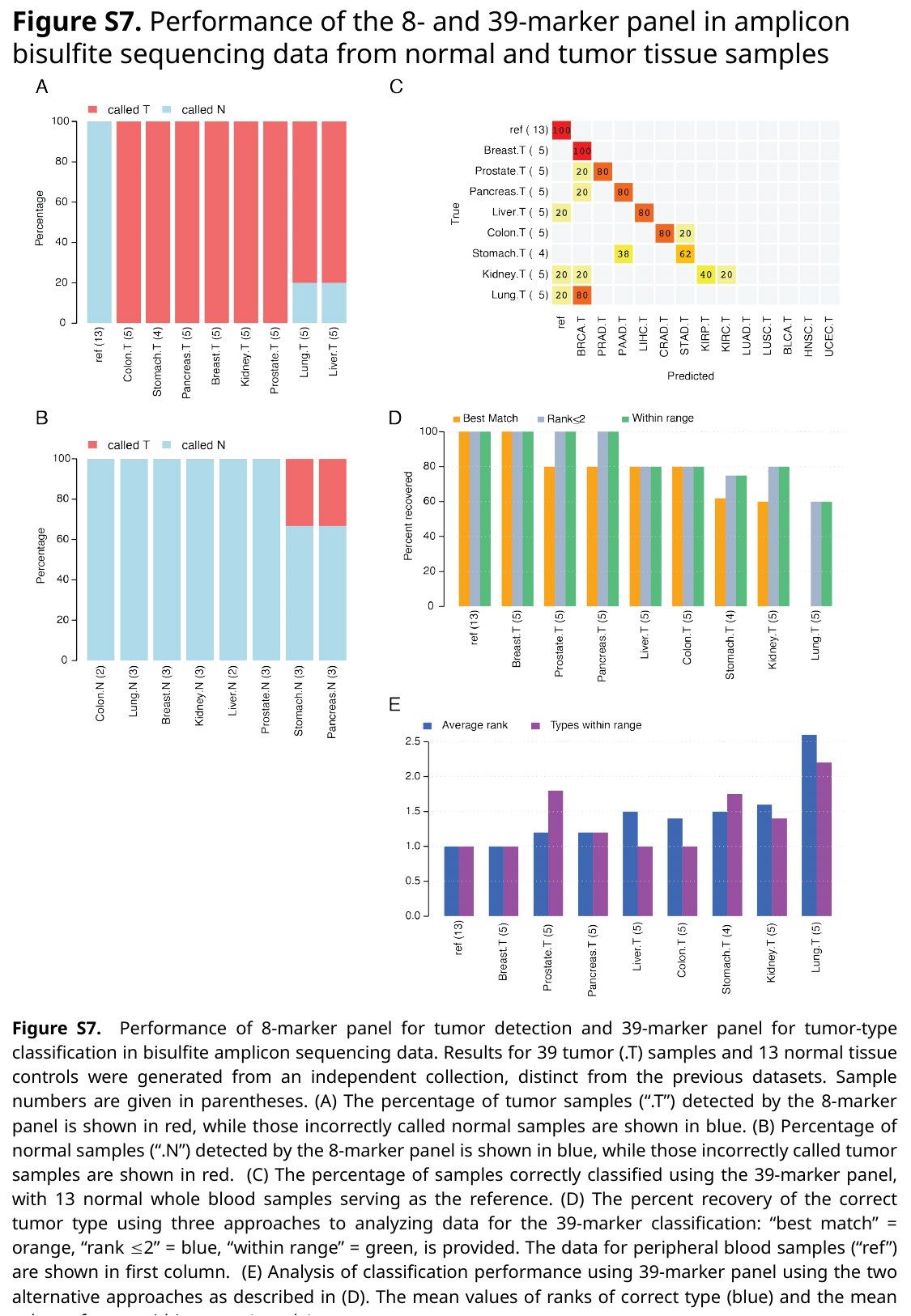

Figure S7. Performance of the 8- and 39-marker panel in amplicon bisulfite sequencing data from normal and tumor tissue samples
Figure S7. Performance of 8-marker panel for tumor detection and 39-marker panel for tumor-type classification in bisulfite amplicon sequencing data. Results for 39 tumor (.T) samples and 13 normal tissue controls were generated from an independent collection, distinct from the previous datasets. Sample numbers are given in parentheses. (A) The percentage of tumor samples (“.T”) detected by the 8-marker panel is shown in red, while those incorrectly called normal samples are shown in blue. (B) Percentage of normal samples (“.N”) detected by the 8-marker panel is shown in blue, while those incorrectly called tumor samples are shown in red. (C) The percentage of samples correctly classified using the 39-marker panel, with 13 normal whole blood samples serving as the reference. (D) The percent recovery of the correct tumor type using three approaches to analyzing data for the 39-marker classification: “best match” = orange, “rank ≤2” = blue, “within range” = green, is provided. The data for peripheral blood samples (“ref”) are shown in first column. (E) Analysis of classification performance using 39-marker panel using the two alternative approaches as described in (D). The mean values of ranks of correct type (blue) and the mean values of types within range (purple).

### Slide 9
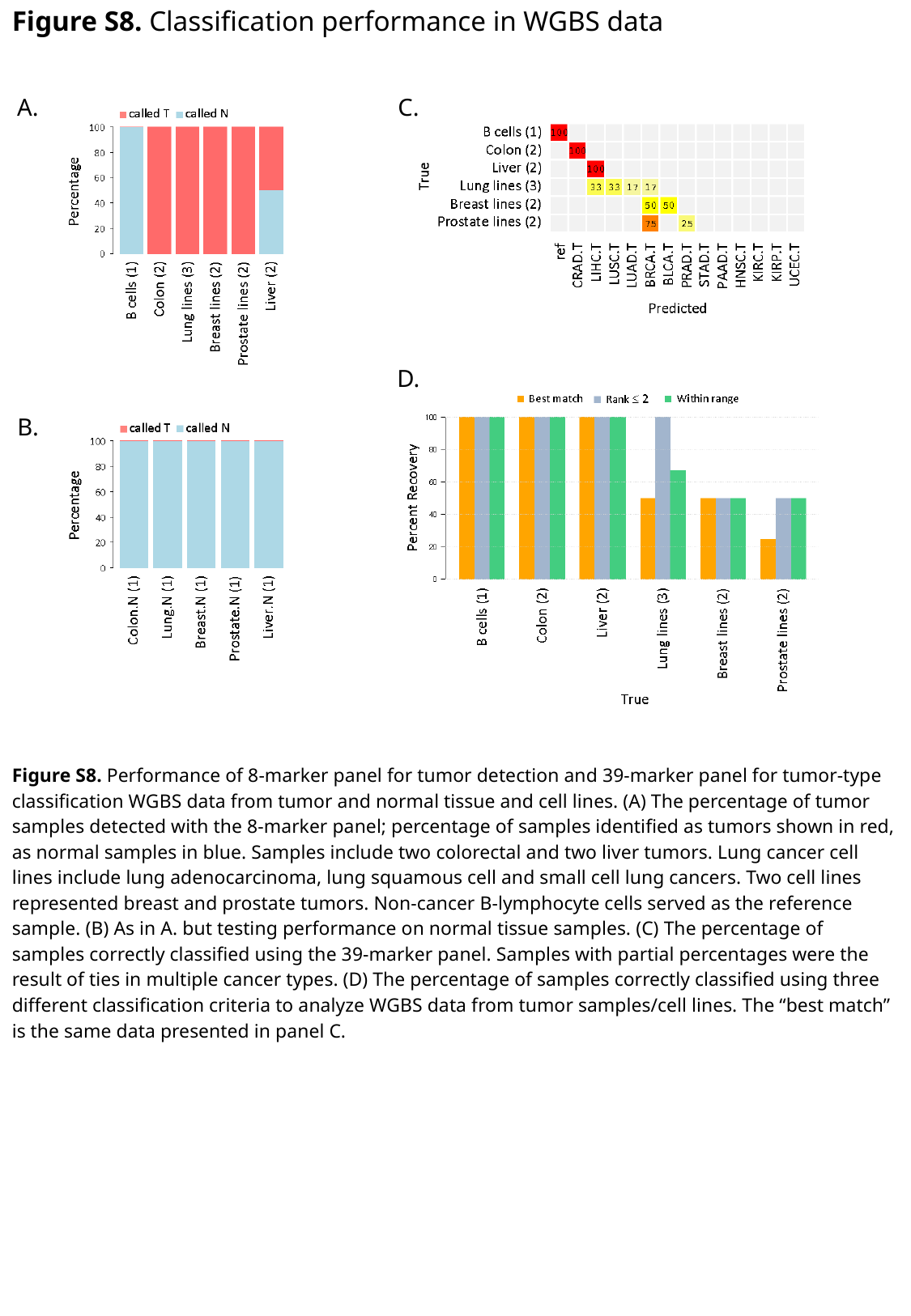

Figure S8. Classification performance in WGBS data
A.
C.
D.
B.
Figure S8. Performance of 8-marker panel for tumor detection and 39-marker panel for tumor-type classification WGBS data from tumor and normal tissue and cell lines. (A) The percentage of tumor samples detected with the 8-marker panel; percentage of samples identified as tumors shown in red, as normal samples in blue. Samples include two colorectal and two liver tumors. Lung cancer cell lines include lung adenocarcinoma, lung squamous cell and small cell lung cancers. Two cell lines represented breast and prostate tumors. Non-cancer B-lymphocyte cells served as the reference sample. (B) As in A. but testing performance on normal tissue samples. (C) The percentage of samples correctly classified using the 39-marker panel. Samples with partial percentages were the result of ties in multiple cancer types. (D) The percentage of samples correctly classified using three different classification criteria to analyze WGBS data from tumor samples/cell lines. The “best match” is the same data presented in panel C.

### Slide 10
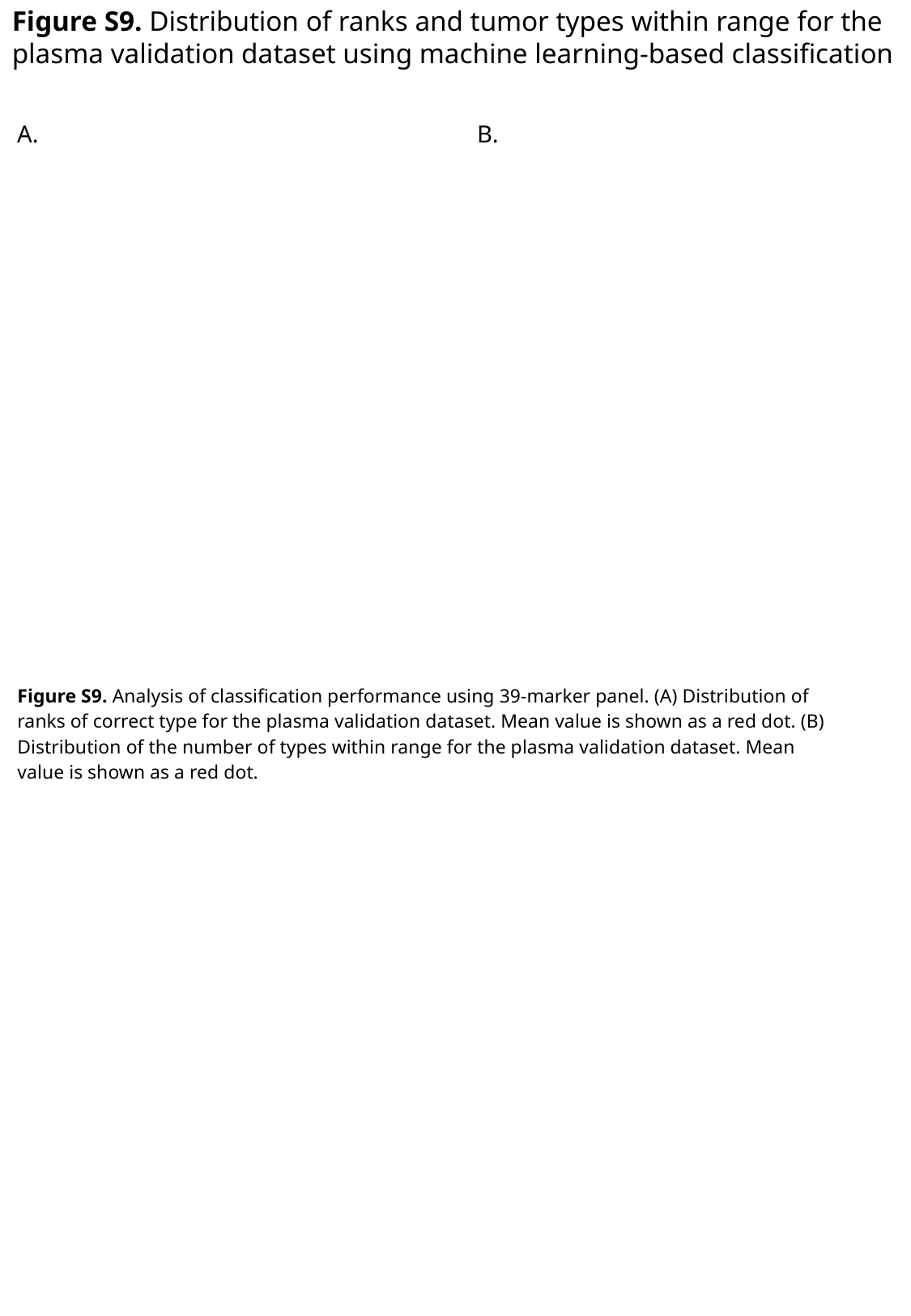

Figure S9. Distribution of ranks and tumor types within range for the plasma validation dataset using machine learning-based classification
A.
B.
Figure S9. Analysis of classification performance using 39-marker panel. (A) Distribution of ranks of correct type for the plasma validation dataset. Mean value is shown as a red dot. (B) Distribution of the number of types within range for the plasma validation dataset. Mean value is shown as a red dot.
