## Supplemental Methods for "Discovery and performance of DNA methylation panels for cancer detection and classification in blood"

### Additional file 3: Supplementary Methods

**Marker selection strategy**

In blood-based cancer diagnostics, we expect the tumor signal to be diluted by the background signal from normal blood cells. Thus, in selecting markers, we looked at loci whose normal methylation represented the extremes of the scale, i.e., virtually absent or saturated, in normal reference samples, with minimal variability. At these loci, even a weakly abnormal signal becomes apparent against the background. More specifically we setup two requirements when screening for markers:

1. Each locus had to have at least four CpGs within 50bp of the probe (including itself)
2. Each probe had to be predominantly unmethylated or methylated in blood reference (median beta value < 0.15 or > 0.85, with standard deviation < 0.15), with less than 3.7% missing data (100 values out of 2,711 samples)

Selection of tumor-normal calling markers

To make a pool of candidate tumor-normal (T-N) markers, we extended the criteria listed above by demanding that each marker had to differ substantially from the blood reference in at least one tumor type (median methylation beta values of tumor samples of that type >0.4 when reference methylation was near zero, or median <0.6 when reference methylation was near 1). There were no conditions imposed on the remaining tumor types; however, median methylation was set to NA for any tumor type (and its normal counterpart) violating the same thresholds as were valid for the reference. Lastly, we considered normal samples from TCGA data and required that all normal tissue types were similar to the reference (normal samples of each of the 13 types satisfied the same thresholds as the blood reference, allowing up to 50% missing values per type). This yielded 4,287 candidate markers, with accompanying median methylation values, or NAs. From this pool of candidate markers, we proceeded by starting with a list of all 13 tumor types, which initially remain unresolved. At each iteration, we chose a probe whose median methylation was substantially different from median methylation in blood reference samples (as defined above), in the maximal number of remaining tumor types (in case of multiple such probes we chose the first one with the maximal absolute difference between the mean methylation across the remaining types in tumors (excluding NAs) and the reference). After the probe was selected, tumor types that were substantially different from reference, were thus resolved and removed from the list of remaining tumor types. Then the chosen marker, as well as all of its neighbors within 100 bp were excluded from subsequent iterations. Iterations proceeded until no tumor types remained, or until the approach failed to find suitable markers. This algorithm can be run multiple times and will select new markers when previous selections (and their neighbors) are excluded from consideration. We chose to use markers from two runs, with each set of markers resolving all of the types, and together the two sets yielded 8 T-N call probes. We call them “unconditional” T-N call probes, because within the collection of 13 tumor types we consider here, these probes can be used to differentiate tumors of all types from normal samples and peripheral blood, without knowing the type a priori.

Sample calling

For each sample, methylation beta values were binarized at the T-N markers. Here, the thresholds for setting the value to 1 were >0.4 for markers with low reference methylation, and <0.6 for markers with high reference methylation, in agreement with probe selection thresholds. A sample was called as tumor if at least one of the binarized values was 1; otherwise, it was called as normal.

Selection of tumor-type classification probes

To preselect a pool of candidate probes to assign tumor samples one of the TCGA types (or blood reference) we employed the two first criteria listed above and additionally added that each probe should have a substantially different methylation level from the blood reference in at least one tumor type (i.e., median > 0.35 when reference methylation was near zero, or median < 0.65 when reference methylation was near 1.0), while the probe’s methylation in all remaining tumor types had to satisfy the same thresholds as were valid for the reference. This resulted in 2,130 candidate probes. We then applied additional filtering, keeping only probes whose target CpGs had average methylation level of either < 0.1 or > 0.9 in the WGBS sequencing of 32 control blood plasma samples reported by Chan et al (1). This reduced the candidate pool to 1,220 loci. From this pool of candidate probes we proceeded to select tumor-type classification markers. First, median beta values of candidate loci within each tumor type and reference were binarized according to below scheme:

|  | **Blood reference** | **TCGA tumor data** | | |
| --- | --- | --- | --- | --- |
| **Median beta value** | <0.15 | >0.35 | >0.15 and ≤0.35 | ≤0.15 |
| **Binarized value** | 0 | 1 | NA | 0 |

|  | **Blood reference** | **TCGA tumor data** | | |
| --- | --- | --- | --- | --- |
| **Median beta value** | >0.85 | <0.65 | ≥0.65 and <0.85 | ≥0.85 |
| **Binarized value** | 0 | 1 | NA | 0 |

In this way, the binarized reference values all end up being set to 0, both for probes with low and with high methylation. Values of 1 in other classification types indicate that the methylation is sufficiently far from reference (as defined by the thresholds), while values of 0 indicate methylation similar to the reference.

We then used binarized candidate marker values to iteratively choose our set of tumor-type classification markers. In each step, we selected a marker with maximal entropy across all subsets of classification types, as described in the following. We call a subset of classification types ambiguous if it has more than one type. We start with an initial single (sub)set of all types together (14 classification types: Peripheral blood, CRAD.T, STAD.T, PAAD.T, BLCA.T, HNSC.T, LUSC.T, LUAD.T, BRCA.T, KIRC.T, KIRP.T, LIHC.T, PRAD.T and UCEC.T). In each iteration step, for each probe/marker in each (ambiguous) subset, the entropy is calculated as $-n\sum_{i=0}^{1} p_{i}\text{log}p_{i}$, where $p_{i}$ are the fractions of *i*’s (0’s and 1’s) in the binarized probe values and *n* is the subset cardinality. In the case of NAs in a subset, the entropy is set to zero. The entropies across the subsets are then added up for each marker, with the intent of choosing a marker with the maximal sum. When there are multiple markers with an identical entropy sum, we choose the first one with smallest Euclidean distance between its median beta values and its binarized values (or their reciprocal, i.e., one minus binarized values), across the types in ambiguous subsets. Given the marker, the subsets in which it has both 0’s and 1’s (and no NAs) are split in two per these values, and this marker is excluded from subsequent iterations. If, after splitting, a new subset contains single classification type, this subset (or type) is no longer ambiguous and is excluded from further iterations. If there are no probes with positive entropy the process stops due to failure; whereas the process stops successfully if there are no ambiguous subsets left to split.

This algorithm for selecting classification markers can be run multiple times, by excluding previously selected markers and possibly also markers in the genomic neighborhood, to obtain new sets of classification markers. We initially compiled three sets of markers (each successfully splitting the classification types) and excluded candidates within 100bp of each selected marker. The three sets together yielded 27 markers. When we performed sample type classification (see next section) using these 27 markers, some tumor types were predicted worse than others. To improve our ability to assign samples to the correct type, we applied the selection algorithm separately to each of the two worst clusters of BLCA-HNSC-LUSC and CRAD-STAD-PAAD tumors, ignoring all other tumor types except the reference. We additionally required each marker to have at least one tumor type satisfying the thresholds attained for the reference (because our goal here was to distinguish between the types, not distinguish between tumor types and reference). This yielded 273 and 1,684 candidate loci, respectively. After triple runs of the algorithm, an additional six probes were added for each cluster, raising the total number of classification probes to 39. The methylation beta values and binarized values of the added probes were set to NA for the types ignored in their selection (Supplementary Figure 1, Additional File 1). One of the 39 probes was also present among the 8 tumor-normal calling probes, thus giving a total of 46 unique probes.

Sample type classification calling

The central classification measurement for each sample we describe is mean distance, defined as follows. Each classification type is represented by values of 39 classification markers, and each sample has methylation beta values at those positions (since WGBS is not deep, we averaged the methylation signal within 50 bases from the probe coordinate). Here we use binarized median beta values for the classification types, as well as binarized beta values of individual samples (using the same thresholds as during marker selection, see scheme in previous section). Note that binarization of individual samples can introduce NAs even when the corresponding classification type marker values are available (i.e., 0 or 1). We use the arithmetic mean (instead of sum) of all non-NA squared differences between the sample and the classification marker values, in line with Euclidean distance calculation. Taking the mean of non-NA values compensates for the possible difference in number of NAs in different samples.

Thus, for each sample we had a vector of mean distances to the classification types. The simplest classification is to the closest type (for most samples, the best predicted type is unique; in case of tied types, sample contribution is split uniformly among the tied types yielding expected classification performance with randomly resolved ties).

**Alternative classification criteria**

From a practical perspective, it would be desirable to establish criteria for how reliable the classification results are for each individual sample and when to consider the second-best and other possibilities. To this end, for each sample we consider several statistics. To estimate our classification performance on the samples of known type we additionally (1) checked whether any of the classes ranked ≤2 (i.e., best or second-best, with nuances in case of ties) were correct, and recorded the rank of the correct class, and (2) defined ranges within which the class measures could be accepted (irrespective of rank), checked whether the correct class was within range, and recorded the number of classes within range. The range was defined as follows: distance up to 1.1*max{shortest distance, 0.1} in case of Euclidian distance measurement. Furthermore, we also made adjustments to the distance measurements itself as described below.

**Comparison of performance by different classification metrics**

In our endeavor to classify tumor types using our 39-marker panel, we encountered nuances in the distances between individual samples and their respective classification types. These variations across different tumor types prompted us to explore enhancements to our standard classification approach. For example, in using our 39-marker panel to classify the tumor type of samples, we relied on ordering distances between individual samples and classification types, or classes (Supplementary figure 2 and 3, Additional file 1). However, distances between individual samples of a given cancer type to their correct class representation differed across types – in some types the samples were typically closer to the expected type than in others. Thus, we considered several modifications to our standard classification approach to compensate for these differences. Specifically, we examined the performance of the 39-marker panel on the TCGA discovery dataset using different approaches to measuring distance. This included the utilization of QFfit (Quantile fraction fit) and variations of a naïve Bayes classifier (Bayes and Bayesf). Our objective was to mitigate the impact of these differences and enhance the accuracy of our classification model. We conclude that the original distance metric (Euclidian distance based) showed the best overall classification performance. It performed slightly better than the Bayes methods, as judged by the percent of samples of each type that were correctly classified using the “best match” approach, as well as the alternate classification criteria “rank≤2” and “within range” (Supplementary figure 4, Additional file 1). To test the alternative criterion (“within range”) we defined ranges for the QFfit method as quantile fraction up to 0.9 and up to 0.95 (i.e., 90% and 95%), and for Bayes classifier we used posterior probability ≥0.1.

Quantile fraction fit

In the first approach, we calculated each sample’s quantile fraction in each classification type, i.e., the fraction of samples of that type with distances less than the one observed (plus half the fraction of samples having distances exactly equal to the one considered). The result for each sample was a vector of sample quantile fractions in the classification types, and the simplest classification would be a type with the smallest fraction. For these calculations (which are calculations of empirical cumulative distribution functions), we used both the raw distributions (i.e., collections of calculated distance values), as well as their parametrizations (see fitted distributions section, below).

Naïve Bayes classifier

In the second approach, naïve Bayes approximation was used to estimate posterior probabilities for a sample to belong to each of considered 14 (classification) types, given the vector of distances. Using Bayes’ formula, this probability is given by

$$P\left( i | \left\{ d_{j} \right\} \right)=\frac{P\left( \left\{ d_{j} \right\} | i \right)P(i)}{\sum_{k} P\left( \left\{ d_{j} \right\} | k \right)P(k)},$$

where $\{d_{j}\}$ is a vector of distances of a given sample to all the classification types, *i* is the considered/possible resulting type in sample classification, and *k* runs through all possible classification types. $P(i)$ are prior probabilities and are taken to be identical in this work; however, they can be adjusted to the observed prevalence of different types of tumor. Naïve Bayes approach approximates $P\left( \left\{ d_{j} \right\} | k \right)$ with $\prod_{j} P(d_{j}|k)$; hence we only need to know the univariate distributions of distances from samples of type *k* to classification type *j*, for all possible values of *j* and *k*. We fit the raw (observed) distributions as described below, and calculated $P(d_{j}|k)$ by integrating the fitted densities in a small interval (0.01) around $d_{j}$. In most cases, one perhaps could use densities instead of probabilities, as Bayes’ formula only contains ratios; however, for an exact zero distance, integration will always yield a non-zero value (even without the point mass at 0, as discussed in the next section), thus allowing for a non-zero estimate for any valid $P\left( \left\{ d_{j} \right\} | k \right)$. The result of this approach is then a vector of estimated posterior probabilities of possible sample types, and the simplest classification would be a type with highest probability.

Fitted distributions of sample distances to classification types

Raw distributions of sample distances to classification types were approximated by beta distributions, with a modification. Due to binarization, there is a noticeable number of exact zero distances. In order to reflect that, we allowed finite masses at the extreme distance values of 0 and 1 (the point mass at 1 was added for symmetry, and its actual mass always was 0). After assigning the distances of exact 0 and 1 to the point masses, the remainder were fit using R function *fitdistr* from package *MASS*. Optionally, we also combined the fitted distributions of the TCGA normal tissue types (.N) together with blood reference distributions, by equally weighting them and using the law of total variance (the point masses at 0 and 1 were not used here). This option was used when calculating fitted quantile fractions, resulting in a substantial increase in the proportion of normal TCGA samples classified as reference, with a smaller effect on TCGA tumors (Supplementary figure 4, Additional file 1, classification using QFfit).

**Addressing overfitting in TCGA and PB_ref_ data**

Generally speaking, designing a classifier and estimating its performance on the same dataset might lead to overfitting (with overoptimistic performance estimates). However, one should not expect noticeable overfitting in our tumor-type classification and T-N calling, as our criteria are primarily based on median methylation values, which are simple, stable and limited summaries of the data. Were we to perform leave-one-sample-out cross-validation, only medians of that sample’s type would be (marginally) affected, if at all, in each round. For an explicit calculation, we split samples of each type into training (~90%) and validation (~10%) sets, and used the training set for probe selection. For the T-N calling, true positive rates were similar in each set (93.7% and 95.1% respectively; both at 95.1% if weighted by number of samples in each type), compared to a lower 90.3% (91.4%, weighted) using the markers derived from all the data and reported in the Results section. However, the false positive rates for TCGA normal tissues were also higher, at 3.6% and 7.5%, respectively (3.6% and 7.6%, weighted), compared with 1.3% (1.2%, weighted) using the markers derived from all of the data. In addition, one of the PB_ref_ training samples (out of 2,440) was miscalled as a tumor. The increased false positive rate in validation samples was due, in large part, to two normal PRAD samples, which were consistently called tumors in multiple scenarios and coincidently ended up in the validation set. In tumor-type classification, training and validation sets yielded 84.6% and 84.0% (weighted 85.1% and 84.9%) correct, respectively (best by distance), compared to 85.3% (weighted 86.1%) using the probes derived from all the data and reported in the Results section. None of the training and two of the validation set PB_ref_ samples were classified incorrectly (as PAAD.T); however, combination of T-N calling and tumor-type classification leads to correct prediction for all reference samples. We conclude that type classification and T-N calling performances are comparable between the training and validation sets.

**Bisulfite amplicon sequencing**

Samples for bisulfite amplicon sequencing

To further validate the markers in our panels, we tested their performance using bisulfite amplicon sequencing. To do so, DNA was obtained from two sources. First, 13 DNA samples from normal blood plasma (1 μg DNA each) were purchased from Fox Chase Cancer Center. This DNA was extracted at Fox Chase Cancer Center using Qiagen Mini-prep kits or the Qiagen Autopure (Germantown, MD). Second, tumor and normal DNA samples (5 μg DNA each) were obtained from Origene from colon, stomach, pancreas, lung, breast, kidney, liver, and prostate tissue samples. There were 5 tumor samples for each tissue, with the exception of 4 samples for stomach. There were 3 normal samples for each tissue, with the exception of 2 samples each for colon and liver. DNA was isolated by Origene (Rockville, MD) using a proprietary protocol similar to the EasyDNA isolation system provided by Invitrogen/ Thermo Fisher Scientific (Waltham, MA).

Bisulfite amplicon sequencing

DNA for bisulfite amplicon sequencing was shipped to Zymo Research for processing and sequencing, as described below. According to their protocol, 500 ng DNA per sample was used for bisulfite conversions. After bisulfite conversion, 50 ng bisulfite-converted DNA was put into the 48-well Fluidigm Access Array system. This translates to roughly 1ng per primer set.

Assays were designed to include preselected CpG sites and multiple neighboring CpGs in the specified regions of interest (ROIs) using primers created with Rosefinch, Zymo Research’s proprietary design tool for sodium-bisulfite-converted-DNA-specific primer generation. We chose parameters such that PCR amplicons would ideally be 100 to 300 bp. In addition, as much as possible, we designed primers that would not anneal to CpG sites in the ROI. In the event that CpG sites were absolutely necessary for target amplification, we ordered primers to be synthesized with a pyrimidine (C or T) at the CpG cytosine in the forward primer or a purine (A or G) in the reverse primer to minimize amplification bias due to either a methylated or unmethylated allele.

All primers arrived in 100 μM TE solution or were resuspended in it. Primers were then mixed (if necessary) and diluted to 2 μM each. Primers were tested using real-time PCR with 1 ng bisulfite-converted control DNA, in duplicate individual reactions. DNA melt analysis was performed to confirm the presence of a specific PCR product. The following guidelines were used to assess performance: (1) had average crossing point (Cp) values <40, (2) duplicate Cps did not have a Cp difference >1 (within 5% coefficient of variation; CV), (3) reached the plateau phase before the run ended at cycle 45, (4) produced melting curves in the expected range for PCR products, and (5) duplicate melts had calculated melting temperatures within 10% CV.

Following primer validation, 500 ng of each DNA sample was bisulfite-converted using the EZ DNA Methylation-Lightning™ Kit (Zymo Research, catalog number D5030) according to the manufacturer’s instructions. We converted 39 tumor samples, including from breast, colon, kidney, liver, lung, pancreas, prostate, and stomach tumors; 22 normal samples, including 2-3 normal samples from each of the previously mentioned organs; and 13 plasma samples from patients without cancer. Amplification of all samples was performed according to the manufacturer’s instructions, using 50 ng bisulfite-converted DNA (roughly 1 ng per primer set), ROI-specific primer pairs, and the Fluidigm Access Array™ System. After barcoding, samples were purified (ZR-96 DNA Clean & Concentrator™, Zymo Research, Cat#D4023) and then prepared for massively parallel sequencing using a MiSeq V2 300bp Reagent Kit and paired-end sequencing protocol, according to the manufacturer’s guidelines. Sequencing read data were aligned using Bismark (2) and extracted as uniquely aligning reads across the amplicons. To calculate methylation at each locus, we used amplicon-wide averages across multiple CpGs. This reduced uncertainty due to data quality concerns.

**Analysis of WGBS data from cell lines and tissue samples**

Whole Genome Bisulfite Sequencing (WGBS) data from derived from Vidal et al. (3). The samples included 11 tumor and tumor cell line samples, 5 normal samples, and 1 B-lymphocyte sample. As mentioned above, since WGBS data is usually not deep, we averaged the methylation signal within 50 bases from the original probe coordinate as used that value similarly to a beta value.

1.	Chan KC, Jiang P, Chan CW, Sun K, Wong J, Hui EP, et al. Noninvasive detection of cancer-associated genome-wide hypomethylation and copy number aberrations by plasma DNA bisulfite sequencing. Proceedings of the National Academy of Sciences of the United States of America. 2013;110:18761-8.

2.	Krueger F, Andrews SR. Bismark: a flexible aligner and methylation caller for Bisulfite-Seq applications. Bioinformatics (Oxford, England). 2011;27:1571-2.

3.	Vidal E, Sayols S, Moran S, Guillaumet-Adkins A, Schroeder MP, Royo R, et al. A DNA methylation map of human cancer at single base-pair resolution. Oncogene. 2017;36:5648-57.
