## Supplementary material for "Discovery and performance of DNA methylation panels for cancer detection and classification in blood": Table S1

### Amount of plasma cfDNA used as input for library preparation

|  | **Mean (ng)** | **Median (ng)** | **Range (ng)** |
| --- | --- | --- | --- |
| **Healthy samples**  **(n = 32)** | 40.0 | 39.9 | 22.3-69.0 |
| **Colon cancer samples (n = 20)** | 117.1 | 40.4 | 23.2-1541.0 |
| **Liver cancer samples (n = 26)** | 165.4 | 60.3 | 20.8-570.4 |
| **Pancreatic cancer**  **(n = 24)** | 136.9 | 46.4 | 20.0-1462.8 |
| **Prostate cancer**  **(n = 23)** | 58.2 | 35.6 | 20.3-570.4 |
| **Stomach cancer**  **(n = 21)** | 149.9 | 47.8 | 23.0-584.2 |
