## Supplementary material for "Discovery and performance of DNA methylation panels for cancer detection and classification in blood": Table S2

### 8 and 39- marker panel probe regions

| **Marker panel** | **Query CpG** | **Illumina ID** | **Twist region** | **target region length** | **target region CpG count** |
| --- | --- | --- | --- | --- | --- |
| pancan8 | chr6:88876741 | cg14186641 | chr6:88876682-88876802 | 121 | 20 |
| pancan8 | chr6:150286508 | cg11660826 | chr6:150286449-150286568 | 120 | 7 |
| pancan8 | chr7:19157193 | cg17839237 | chr7:19157134-19157253 | 120 | 10 |
| pancan8 | chr10:14816201 | cg13840308 | chr10:14816142-14816261 | 120 | 6 |
| pancan8 | chr12:129822259 | cg01163404 | chr12:129822200-129822319 | 120 | 4 |
| pancan8 | chr14:89628169 | cg21898844 | chr14:89628110-89628229 | 120 | 5 |
| pancan8 | chr17:40333009 | cg17812788 | chr17:40332950-40333069 | 120 | 16 |
| pancan8 | chr17:46655394 | cg14458834 | chr17:46655335-46655454 | 120 | 16 |
| tumortype39 | chr2:114035619 | cg09250933 | chr2:114035560-114035679 | 120 | 6 |
| tumortype39 | chr2:176994448 | cg14473102 | chr2:176994389-176994508 | 120 | 19 |
| tumortype39 | chr2:176994764 | cg24416513 | chr2:176994705-176994824 | 120 | 18 |
| tumortype39 | chr2:219256101 | cg16926316 | chr2:219256042-219256161 | 120 | 8 |
| tumortype39 | chr2:240270793 | cg04869380 | chr2:240270734-240270853 | 120 | 4 |
| tumortype39 | chr2:25600752 | cg17774001 | chr2:25600693-25600812 | 120 | 5 |
| tumortype39 | chr2:61372138 | cg16328106 | chr2:61372079-61372198 | 120 | 9 |
| tumortype39 | chr2:66665428 | cg00995986 | chr2:66665369-66665488 | 120 | 8 |
| tumortype39 | chr2:8724060 | cg18091469 | chr2:8724001-8724120 | 120 | 9 |
| tumortype39 | chr4:142054417 | cg03563667 | chr4:142054358-142054477 | 120 | 14 |
| tumortype39 | chr4:156588387 | cg23912454 | chr4:156588328-156588447 | 120 | 11 |
| tumortype39 | chr5:140306231 | cg23026864 | chr5:140306172-140306291 | 120 | 12 |
| tumortype39 | chr6:106958645 | cg22231056 | chr6:106958586-106958705 | 120 | 11 |
| tumortype39 | chr6:133562470 | cg11942956 | chr6:133562411-133562530 | 120 | 18 |
| tumortype39 | chr7:27196759 | cg08934785 | chr7:27196700-27196819 | 120 | 6 |
| tumortype39 | chr7:4801993 | cg23871697 | chr7:4801934-4802053 | 120 | 5 |
| tumortype39 | chr8:1895558 | cg20690085 | chr8:1895499-1895618 | 120 | 6 |
| tumortype39 | chr8:97506675 | cg04261408 | chr8:97506616-97506735 | 120 | 7 |
| tumortype39 | chr8:102451058 | cg19882915 | chr8:102450998-102451118 | 121 | 6 |
| tumortype39 | chr9:140683797 | cg14050824 | chr9:140683738-140683857 | 120 | 6 |
| tumortype39 | chr10:103603810 | cg14489801 | chr10:103603751-103603870 | 120 | 6 |
| tumortype39 | chr10:1120831 | cg11314310 | chr10:1120772-1120891 | 120 | 11 |
| tumortype39 | chr10:114591733 | cg02571204 | chr10:114591673-114591793 | 121 | 10 |
| tumortype39 | chr10:116064472 | cg24507921 | chr10:116064413-116064532 | 120 | 5 |
| tumortype39 | chr10:21788638 | cg18621142 | chr10:21788579-21788698 | 120 | 7 |
| tumortype39 | chr10:5566908 | cg24831879 | chr10:5566849-5566968 | 120 | 8 |
| tumortype39 | chr10:8097331 | cg14327531 | chr10:8097272-8097391 | 120 | 8 |
| tumortype39 | chr10:8097689 | cg15267232 | chr10:8097629-8097750 | 122 | 15 |
| tumortype39 | chr11:60619955 | cg12528056 | chr11:60619896-60620015 | 120 | 13 |
| tumortype39 | chr11:8284312 | cg26121782 | chr11:8284253-8284372 | 120 | 9 |
| tumortype39 | chr12:54427173 | cg17031478 | chr12:54427114-54427233 | 120 | 17 |
| tumortype39 | chr13:113424938 | cg02699898 | chr13:113424879-113424998 | 120 | 9 |
| tumortype39 | chr16:51184392 | cg00582524 | chr16:51184333-51184452 | 120 | 11 |
| tumortype39 | chr16:678127 | cg06013113 | chr16:678068-678187 | 120 | 8 |
| tumortype39 | chr17:46655394 | cg14458834 | chr17:46655335-46655454 | 120 | 16 |
| tumortype39 | chr17:46711341 | cg01452847 | chr17:46711281-46711401 | 121 | 16 |
| tumortype39 | chr19:16189360 | cg06590173 | chr19:16189301-16189420 | 120 | 4 |
| tumortype39 | chr19:1827498 | cg18751958 | chr19:1827439-1827558 | 120 | 8 |
| tumortype39 | chr19:18335182 | cg18816098 | chr19:18335123-18335242 | 120 | 10 |
