## Supplementary material for "Discovery and performance of DNA methylation panels for cancer detection and classification in blood": Table S3

### Probe status for the 8-marker panel as defined by median-based cutoff in methylation array data

| **Probe/Tumor** | cg14186641 | cg11660826 | cg17839237 | cg13840308 | cg01163404 | cg21898844 | cg17812788 | cg14458834 |
| --- | --- | --- | --- | --- | --- | --- | --- | --- |
| BLCA |  | Hyper |  |  | Hypo |  |  |  |
| BRCA |  | Hyper |  |  | Hypo |  | Hyper |  |
| COAD |  | Hyper | Hyper |  | Hypo |  |  |  |
| HNSC |  |  |  |  | Hypo |  |  | Hyper |
| KIRC |  |  |  | Hypo |  | Hypo |  |  |
| KIRP |  |  |  | Hypo |  | Hypo |  |  |
| LIHC |  | Hyper |  |  | Hypo |  | Hyper |  |
| LUAD |  |  |  |  |  |  | Hyper | Hyper |
| LUSC |  |  |  |  | Hypo |  |  | Hyper |
| PAAD | Hyper |  | Hyper |  |  |  |  |  |
| PRAD |  | Hyper |  |  |  |  | Hyper |  |
| READ |  | Hyper | Hyper |  | Hypo |  |  |  |
| STAD | Hyper | Hyper | Hyper |  | Hypo |  |  | Hyper |
| UCEC |  | Hyper |  |  | Hypo |  |  |  |
