## Supplementary material for "Discovery and performance of DNA methylation panels for cancer detection and classification in blood": Table S4

### Table S4: Additional clinical information on patient cohorts

|  | **Colon**  **cancer** |
| --- | --- |
| **No. of patients** | 20 |
| **Age**, median (range) | 60 (33-82) |
| **Sex**, n(%) |  |
| Male | 17 (85%) |
| Female | 3 (15%) |
| **Pathologic stage group** |  |
| IIA | 4 |
| IIB | 1 |
| IIIA | 1 |
| IIIB | 10 |
| IIIC | 2 |
| IV | 1 |
| NA | 1 |
| **Histology** |  |
| Adenocarcinoma | 20 |
| **Tumor site** |  |
| Colon, sigmoid | 9 |
| Colon, ascending | 3 |
| Colon, descending | 1 |
| Colon, transverse | 1 |
| Colon, splenic flexure | 1 |
| Cecum | 3 |
| Appendix | 1 |
| Colon (not specified) | 1 |

|  | **Liver**  **cancer** |
| --- | --- |
| **No. of patients** | 26 |
| **Age**, median (range) | 65 (40-74) |
| **Sex**, n(%) |  |
| Male | 10 (58%) |
| Female | 6 (42%) |
| **Pathologic stage group** |  |
| IB | 1 |
| I | 4 |
| IIB | 1 |
| II | 3 |
| IIIA | 2 |
| IVA | 1 |
| IVB | 1 |
| IV | 2 |
| NA | 11 |
| **Histology** |  |
| Hepatocellular carcinoma | 17 |
| Adenocarcinoma | 1 |
| Cholangiocarcinoma | 6 |
| Leiomyosarcoma | 2* |
| **Tumor site** |  |
| Liver | 26 |

** The same patient donated two blood samples, 3 years apart. Pathologic stage information was unavailable at both time points and no information about progression of disease was recorded. For this study, the samples were regarded as independent samples.*

|  | **Pancreatic**  **cancer** |
| --- | --- |
| **No. of patients** | 24 |
| **Age**, median (range) | 60 (38-78) |
| **Sex**, n(%) |  |
| Male | 15 (62.5%) |
| Female | 9 (37.5%) |
| **Pathologic stage group** |  |
| IB | 1 |
| IIA | 2 |
| IIB | 7 |
| III | 1 |
| IVA | 1 |
| IV | 11 |
| NA | 1 |
| **Histology** |  |
| Adenocarcinoma | 12 |
| Carcinoid tumor | 2 |
| Carcinoma | 1 |
| Infiltrating duct carcinoma | 6 |
| Invasive carcinoma | 1 |
| Neuroendocrine carcinoma | 2 |
| **Tumor site** |  |
| Body of pancreas | 1 |
| Head of pancreas | 18 |
| Tail of pancreas | 5 |

|  | **Prostate**  **cancer** |
| --- | --- |
| **No. of patients** | 23 |
| **Age**, median (range) | 61 (49-75) |
| **Sex**, n(%) |  |
| Male | 23 (100%) |
| Female | - |
| **Pathologic stage group** |  |
| IIC | 1 |
| IIIB | 7 |
| IIIC | 3 |
| III | 2 |
| IVA | 2 |
| NA | 1 |
| **Grade group** |  |
| 1 | 1 |
| 2 | 4 |
| 3 | 1 |
| 4 | 1 |
| **Histology** |  |
| Adenocarcinoma | 23 |
| **Tumor site** |  |
| Prostate gland | 23 |

|  | **Stomach**  **cancer** |
| --- | --- |
| **No. of patients** | 21 |
| **Age**, median (range) | 66 (38-80) |
| **Sex**, n(%) |  |
| Male | 16 (76%) |
| Female | 5 (24%) |
| **Pathologic stage group** |  |
| IIA | 3 |
| IIB | 5 |
| IIIA | 2 |
| III | 4 |
| IVB | 6 |
| NA | 1 |
| **Histology** |  |
| Adenocarcinoma | 17 |
| Discohesive carcinoma | 1 |
| Mucinous adenocarcinoma | 2 |
| Tubular adenocarcinoma | 1 |
